# The human Dicer helicase domain mediates ATP hydrolysis and RNA rearrangement

**DOI:** 10.1101/2023.06.28.546842

**Authors:** Kinga Ciechanowska, Agnieszka Szczepanska, Kamil Szpotkowski, Klaudia Wojcik, Anna Urbanowicz, Anna Kurzynska-Kokorniak

## Abstract

Vertebrates have one Dicer ortholog that generates both microRNAs (miRNAs) and small interfering RNAs (siRNAs), in contrast to the multiple Dicer-like proteins found in flies and plants. Here, we focus on the functions of the human Dicer (hDicer) helicase domain. The helicase domain of hDicer is known to recognize pre-miRNA substrates through interactions with their apical loop regions. Besides interacting with canonical substrates, the hDicer helicase domain has also been suggested to bind many different cellular RNAs; however, a comprehensive study of the biochemical activities and substrate specificity of the hDicer helicase domain towards different nucleic acids has yet to be undertaken. Here, we conduct such an analysis and reveal that full-length hDicer, through its helicase domain, hydrolyzes ATP. We also show that the hDicer helicase domain binds single-but not double-stranded RNAs and DNAs and that a structural rearrangement of the substrate accompanies the binding of single-stranded RNAs. This RNA rearrangement activity is ATP-independent. Our findings open new avenues for future studies aimed at defining the cellular activities of hDicer that may be associated with these newly described biochemical properties.

## 1. Introduction

Dicer ribonucleases are the members of the ribonuclease III (RNase III) family, which function as double-stranded RNA (dsRNA) specific endoribonucleases. In mammals, only one gene encoding the Dicer protein has been identified [1], while some other organisms encode multiple Dicers, with different versions specialized for miRNA or siRNA production [2-5]. Human Dicer (hDicer) consists of 1,922 amino acids (∼220 kDa) and comprises an amino (N)-terminal helicase domain, a domain of unknown function (DUF283), a Platform domain, Piwi-Argonaute-Zwille (PAZ) domain, a Connector helix, two RNase III domains (IIIa and IIIb) and a dsRNA-binding domain (dsRBD) (**Fig. 1a**) [6]. The Platform-PAZ-Connector helix fragment is often called “the PAZ cassette” [7] or “the PPC cassette” [8]. The three-dimensional structure of hDicer resembles the letter L (**Fig. 1b**) [9]. The roles of the individual hDicer domains in binding and processing canonical substrates, i.e., single-stranded hairpin precursors of miRNAs (pre-miRNAs), have been studied extensively. The helicase domain selectively interacts with the apical loop of pre-miRNAs, thus promoting substrate discrimination [10, 11]. The DUF283 domain is implicated in the binding of single-stranded nucleic acids [12], and, therefore, may also be involved in interactions with the apical loop of pre-miRNA hairpins [9]. The PPC cassette anchors the 5’ phosphate and 2-nucleotide (nt) 3’ overhang of the pre-miRNA substrates [7]. The RNase IIIa and RNase IIIb domains form a dsRNA-cleavage center [6]. The carboxy (C)-terminal dsRBD plays a supporting role in pre-miRNA binding [13], and together with the helicase and the DUF283 domains, controls substrate access to the catalytic core of hDicer [11, 14].

**Fig. 1.**
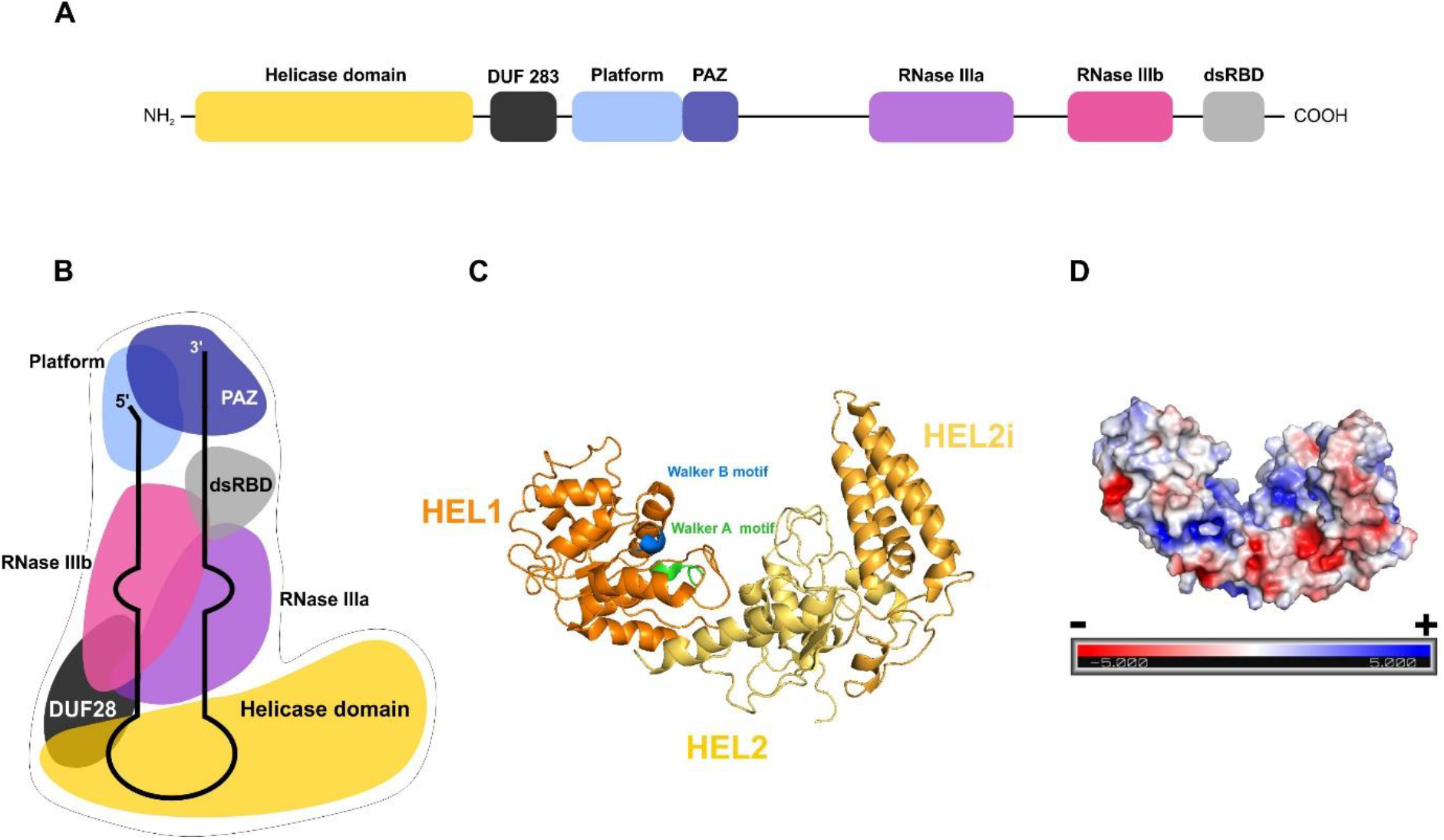
Architecture of human ribonuclease Dicer (hDicer) and its helicase domain. **a** Schematic showing the domain organization of hDicer based on [27]. **b** Schematic of the tertiary structure of hDicer with a miRNA precursor (pre-miRNA) based on [10] and [9]. **c** The tertiary structure of the hDicer helicase domain (PDB entry 5ZAL) visualized using PyMOL. Three subdomains: HEL1, HEL2i and HEL2 are distinguished, and the DExD/H-box including a Walker B motif and Walker A motif are indicated. **d** The distribution of charges on the surface of the helicase domain (PDB entry 5ZAL) visualized using PyMOL.

The helicase domain is believed to be one of the most conserved regions in Dicer [3]; its sequence is highly similar to the SF2 helicases [15-17]. SF2 is the largest superfamily of helicases and translocases, and enzymes belonging to this superfamily are implicated in diverse cellular processes, including transcription, translation, RNA processing, and decay [18]. The SF2 helicases are divided into a number of subfamilies including, but not limited to, the DExD/H-box RNA and DNA helicases and RIG-I-like helicases [19]. The DExD/H-box motif (also called the Walker B motif) is essential for ATP hydrolysis and is present in helicases which unwind structures in dsRNA and double-stranded DNA (dsDNA) [18]. The RIG-I motif is found in ATP-dependent dsRNA translocases [19].

The hDicer helicase domain consists of three subdomains: HEL1, HEL2i and HEL2, which form three lobes in the tertiary structure (**Fig. 1c**) [3]. The HEL1 subdomain contains the DExD/H-box motif, HEL2i includes a RIG-I motif, while the HEL2 subdomain, also called “helicase C”, is conserved in all helicases from the DExD/H and the RIG-I-like families of SF2 helicases [20]. It must be emphasized that not all Dicer proteins contain all three helicase subdomains, and some may not contain the helicase domain at all. For example, the HEL1 subdomain of *Drosophila melanogaster* Dicer-1, which generates miRNAs, has degenerated and is incapable of ATP hydrolysis [4]. Similarly, the Dicer-type protein from the fungus *Magnaporthe oryzae* lacks a HEL2i subdomain [21], whereas the entire helicase domain is missing from *Giardia intestinalis* Dicer [22]. Moreover, the presence of the DExD/H-box and RIG-I motifs does not guarantee that the protein will have the ability to unwind double-stranded nucleic acids or translocate. Examples of such proteins include vertebrate Dicers, for which double-stranded nucleic acid unwinding and translocase activities have not been demonstrated. In addition, there have been no reports on the ATP-hydrolysis activity of the vertebrate Dicer helicase domains thus far [23]. In contrast, there are several examples of invertebrate, fission yeast, and plant Dicer proteins whose helicase domains display ATP-dependent translocation activity; e.g., *D. melanogaster* Dicer-2 [24], *Caenorhabditis elegans* Dicer [24], *Schizosaccharomyces pombe* Dicer [25] and plant Dicer-like proteins (DCL proteins) from *Arabidopsis thaliana* and *Medicago truncatula* [2, 26, 27]. These Dicer-type proteins also have functional RIG-I type subdomains (HEL1, HEL2i and HEL2). Moreover, these Dicer-type proteins act as viral RNA sensors that activate appropriate effector RNA interference (RNAi) pathways and, consequently, trigger viral RNA degradation [28]. It has been proposed that vertebrate Dicer helicase domains lost their ability to recognize dsRNAs because vertebrates have other intracellular RNA virus sensors; for instance, RIG-I helicases [29]. Indeed, vertebrate RIG-I helicases serve as receptors for the innate immune system during infection by RNA viruses [30]. Once stimulated by the viral dsRNA, the RIG-I receptor initiates signaling pathways that trigger interferon-β production and activation of many genes involved in the innate immune response [31]. Consequently, the helicase domains of invertebrate and vertebrate Dicer type-proteins might have developed different functions.

The helicase domain of Dicer proteins can also serve as a binding platform for dsRNA Binding Proteins (dsRBPs); e.g., the hDicer helicase domain can mediate formation of a complex with the TAR RNA-binding protein (TRBP) or the protein activator of protein kinase R (PACT) [32]. TRBP and PACT are important regulators that promote substrate binding during small regulatory RNA production [33]. In addition, it was recently shown that hDicer specifically interacts with several dsRBPs and RNA helicases during viral infection [34]. Specifically, proteins such as DExH-Box Helicase 9 (DHX9), adenosine deaminase acting on RNA 1 (ADAR-1) and protein kinase RNA-activated (PKR) were shown to interact with hDicer in virus-infected cells, and the helicase domain was shown to be essential for interactions with these proteins [34]. Thus, the helicase domain may control the recruitment of different factors that diversify the functions of Dicer proteins.

Growing evidence points to possible cleavage-independent regulatory roles of Dicer. For example, it was demonstrated that in *C. elegans* and human cells, Dicer can bind various RNAs, including mRNAs and long noncoding RNAs (lncRNAs) without further cleavage, so-called “passive binding” [35]. “Passive sites” present within transcripts were proposed to function as a buffering system that sequesters Dicer cleavage activity away from pre-miRNAs [35]. Additionally, Dicer interaction with mRNAs may regulate transcript stability in the cell [35]. Passive binding of hDicer to cellular transcripts is hypothetically mediated by its helicase domain [35]. Indeed, distribution of the charges on the surface of the helicase domain reveals a positively charged groove which can potentially bind nucleic acids (**Fig. 1d**). Nevertheless, a comprehensive characterization of the biochemical activities of the hDicer helicase domain and its substrate specificity towards different nucleic acids has yet to be reported. Here, we present insight into the biochemical properties of the hDicer helicase in the context of different RNA and DNA substrates, both single- and double-stranded.

## 2. Materials and Methods

### 2.1. Oligonucleotides

DNA and RNA oligonucleotides were purchased from Genomed (Warsaw, Poland) and FutureSynthesis (Poznan, Poland), respectively. Sequences of all oligonucleotides used in this study are listed in Table 1.

**Table 1.**
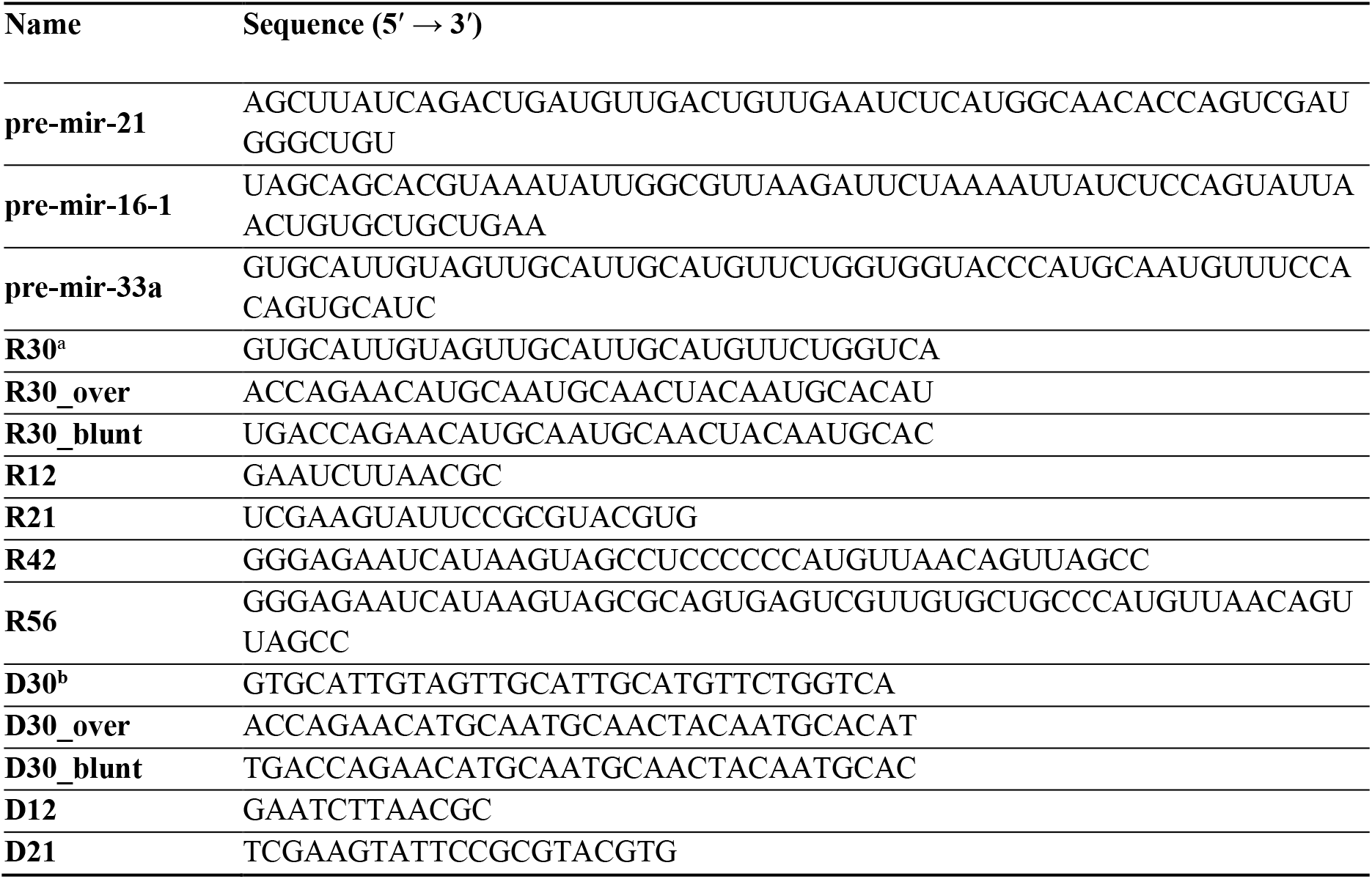
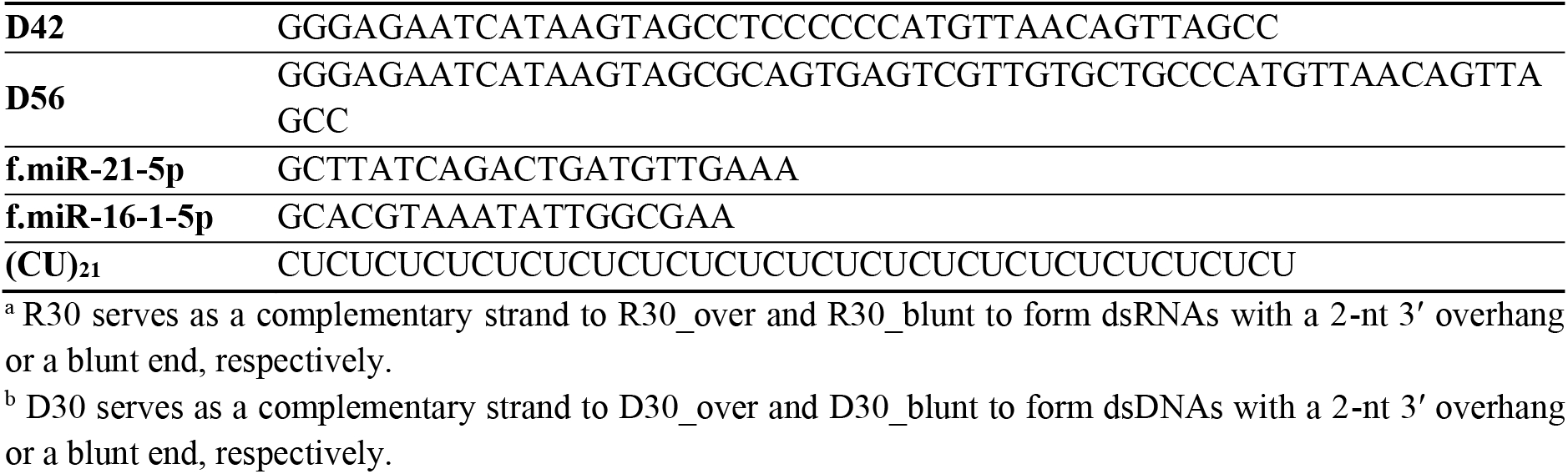
Sequences of the oligonucleotides used in this study.

### 2.2. ^*32*^P labeling of oligonucleotides

The 5′-end labeling of oligonucleotides was performed according to a previously described procedure [36].

### 2.3. Preparation of dsRNA and dsDNA

To prepare dsRNA and dsDNA substrates, non-labeled strand (R30_sense/D30_sense or R30_over/D30_over) was hybridized at a ∼1:1 molar ratio with ^32^P-labeled complementary strand (R30/D30) by heating to 95 °C and then slowly cooling to room temperature in a buffer containing 50 mM NaCl, 2.5 mM MgCl_2_ and 20 mM Tris-HCl, pH 7.5. Next, the reaction mixtures were PAGE-purified using 12% native PAA gels to obtain pure, double-stranded fractions free of single-stranded species.

### 2.4. ATP hydrolysis assay

ATP hydrolysis reactions were carried out in 10-µl volumes. HEL (2 nM) or WT hDicer (2 nM for time-course and 0.36, 0.72, 1.44, 2.88, 5.75, 11.5 nM for protein concentration-dependent hydrolysis) or hDicer_ΔHEL (2 nM) or hDicer_K70A (2, 20, 200 nM for time-course and 36, 72, 144, 288, 575, 1150 nM for protein concentration-dependent hydrolysis) was added to 10,000 cpm (2 nM) [γ^32^P]-ATP (Hartman Analytic, Braunschweig, Germany) and incubated in binding buffer (50 mM NaCl, 150 mM Tris-HCl pH 7.5) with 2.5 mM MgCl_2_ for 1, 5, 15, 30, 60, 90 and 120 min for time-course or 30 min for protein concentration-dependent hydrolysis at 37 °C. Control reactions were prepared without protein. The reactions were separated in 10% native PAA gels at 4 °C in 1x TBE running buffer. The data were collected using an Amersham™ Typhoon™ (Cytiva, Washington, D.C., USA) and quantified using MultiGauge 3.0 (Fujifilm, Tokyo, Japan). The ATP hydrolysis assays were conducted in triplicate.

### 2.5. ATP-binding assay

ATP binding reactions were carried out in 20-µl volumes. WT hDicer (3.6, 7.2, 14.4, 28.8, 57.5, 115 nM) or hDicer_K70A (36, 72, 144, 288, 575, 1150 nM) was added to 10,000 cpm (2 nM) [γ^32^P]-ATP (Hartman Analytic, Braunschweig, Germany) and incubated in binding buffer (50 mM NaCl, 150 mM Tris-HCl pH 7.5) with 2.5 mM MgCl_2_ for 1 min at 4 °C. The reactions were separated in 10% native PAA gels at 4 °C in 1x TBE running buffer. The data were collected using a Amersham™ Typhoon™ (Cytiva, Washington, D.C., USA) and quantified using MultiGauge 3.0 (Fujifilm, Tokyo, Japan). The ATP binding assays were conducted in triplicate.

### 2.6. Binding assay

The reactions were carried out in 40-µl volumes. HEL (2.97, 5.94, 11.86, 23.75, 47.5, 95 µM) was added to 10,000 cpm (2.5 nM) of ^32^P-labeled RNA, DNA, dsRNA or dsDNA and incubated in binding buffer (50 mM NaCl, 150 mM Tris-HCl pH 7.5) for 15 min at room temperature. Control reactions were prepared without protein. The reactions were separated in 5% native PAA gels at 4 °C in 1x TBE running buffer. The data were collected using a Amersham™ Typhoon™ (Cytiva, Washington, D.C, USA) and quantified using MultiGauge 3.0 (Fujifilm, Tokyo, Japan). Binding assays were conducted in triplicate.

### 2.7. Data analysis

Binding assay results were used to estimate the equilibrium dissociation constant (K_d_). K_d_ was estimated on the basis of densitometry analysis in Multi Gauge software (Fujifilm, Tokyo, Japan). K_d_ was calculated using formula:

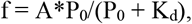

where: f - fraction of bound substrate; P_0_ - molar concentrations of the protein (µM); A - maximum RNA/DNA bound (%). K_d_ was calculated for f = 0.5 (half of the substrate is bound). P_0_represented the protein concentration at which 50% of substrate was bound. K_d_ values were calculated based on results from three experiments.

### 2.8. Bio-Layer Interferometry (BLI)

The measurements were carried out in 200-µl volumes on black 96-well plates (Greiner bio-one, Kremsmünster, Austria) using Octet K2 (ForteBio, Pall Life Sciences, New York, USA) and Octet NTA biosensors (Sartorius, Göttingen, Germany). HEL (1 µM) was incubated with ssRNAs: R20, R30, R40, R50; or ssDNAs: D20, D30, D40, D50 in binding buffer (50 mM NaCl, 150 mM Tris-HCl pH 7.5); increasing amounts of substrate were used: 3.125 µM, 6.25 µM, 12.5 µM, 25 µM, 50 µM, 100 µM. After each measurement, HEL buffer (50 mM HEPES buffer pH 7.5, 0.5 M NaCl, 0.1% Triton X-100 and 5% glycerol), glycine (10 mM, pH 1.7) and NiSO_4_ (10 mM) were used for neutralization and regeneration of Ni-NTA sensor. Measurements were carried out at 23°C at a shaking speed of 1,000 rpm and according to the following steps: Loading (1 800 s), Baseline (60 s), Association (20 s), Dissociation (20 s), Neutralization (3x 30 s), Regeneration (60 s). Each measurement included a parallel reference, in which HEL-loaded biosensors were immersed in the binding buffer lacking ligand. Reference subtracted BLI response curves were generated and used for the determination of the K_d_ constant and its error. Inter-step correction and Y-alignment were used to minimize tip-dependent variability. Data were collected and globally fitted by a 1:1 stoichiometry model using the Data Acquisition and Data Analysis Software vHT 11.1 (ForteBio, Pall Life Sciences, New York, USA). The fitting met the quality criteria χ2 < 3 and R2 ≥ 0.96.

### 2.9. Circular Dichroism (CD)

Circular dichroism spectra were collected on a J-815 CD spectrometer (JASCO, Tokyo, Japan) equipped with a Peltier thermostatic cell holder. HEL (28 µM), RNA (28 µM), and RNA•HEL complexes (molar ratio 1:1) were placed in buffer containing 10 mM HEPES pH 8.0, 300 mM NaF, incubated at 15° C, and analyzed in a 0.1 cm quartz cuvette (Hellma 100-QS, Jena, Germany). Each CD spectrum was generated based on 9 scans in continuous scanning mode, with a scanning speed of 50 nm min^-1^, a 1 nm bandwidth, a 0.5 nm data pitch and a data integration time 1 s. When collecting a regular spectrum, data were gathered at wavelengths ranging from 210 to 350 nm for the thermal melt analysis. Thermal analyses were conducted in a temperature range from 5 to 90 °C. Buffer subtraction and all spectra processing were made using the Jasco Spectra Menager software and Savitzky-Golay tool with a smoothing window of 10 points. The normalized root mean square deviation (NRMSD) for each CD spectrum analysis was less than 0.1. CD data are presented in terms of ellipticity values in millidegrees (mdeg).

### 2.10. Cell culture and transfection

293T NoDice cells [37] were cultured in DMEM (Gibco, Thermo Fisher Scientific, Waltham, MA, USA) supplemented with 10% FBS (Gibco, Thermo Fisher Scientific, Waltham, MA, USA), Penicillin-Streptomycin (100 U/mL of penicillin and 100 μg/mL of streptomycin, Gibco) and 1 mM Sodium Pyruvate (Gibco, Thermo Fisher Scientific, Waltham, MA, USA), as described in Bogerd and colleagues [37]. Transfection of plasmids carrying wild-type hDicer, hDicer_ΔHEL and hDicer_K70A expression plasmids was carried out using the DharmaFECT kb DNA Transfection Reagent (Dharmacon, Lafayette, CO, USA) according to the manufacturer’s instructions. The 293T NoDice cell line was kindly provided by Prof. Bryan R. Cullen [37].

#### 2.1.1. Protein preparations used in the studies

The HEL cDNA (1-624 aa hDicer) was amplified by PCR using a purchased plasmid encoding a complete *Homo sapiens* Dicer1 ribonuclease type III sequence (PubMed, NM_030621) (GeneCopoeia, Rockville, MD, USA). The obtained fragment was cloned into pMCSG7 vector (courtesy of Laboratory of Protein Engineering, Institute of Bioorganic Chemistry, Polish Academy of Sciences), which introduces a His6-tag at the N-terminus of the protein. HEL was expressed in *E. coli* strain BL21Star (Thermo Fisher Scientific, Waltham, MA, USA) in standard Luria-Bertani (LB) medium. *E. coli* cells were treated with 0.4 mM IPTG and cultured for 18 hours at 18 °C with shaking. Cell pellets were then isolated, lysed, and protein purified using Ni^2+^-Sepharose High Performance beads (Cytiva, Washington, D.C., USA) with an imidazole gradient (0.02 M – 1 M) in 0.05 mM HEPES buffer (pH 7.5) supplemented with 0.5 M NaCl, 0.1% Triton X-100 and 5% glycerol. Additional washing with 1 M NaCl solution was applied to remove nucleic acid contamination. Protein purity was assessed by SDS-PAGE. Protein was concentrated using Amicon filters (Merck, Darmstadt, Germany) in buffer (50 mM HEPES pH 7.5, 500 mM NaCl, 0.1% Triton X-100, 5% glycerol) and stored at -80 °C.

The expression plasmid encoding the wild-type hDicer was prepared as described previously [38]. In brief, the expression plasmid was obtained using PCR amplification. All primers were designed based on the cDNA encoding transcript 2 of human *DICER1* (NM_030621.4). Expression plasmids were constructed with the obtained PCR product using the SureVector system (Agilent, Santa Clara, CA, USA) according to the manufacturer’s instructions. The expression plasmid encoding the hDicer_ΔHEL variant was purchased from Addgene (#51366) [39], as was plasmid encoding hDicer_K70A (#41589) [40].

The wild-type hDicer, hDicer_ΔHEL and hDicer_K70A were produced and purified according to a previously described procedure [38]. Obtained hDicer, hDicer_ΔHEL and hDicer_K70A proteins were examined using Western blot analysis according to a previously described procedure [38].

#### 2.1.2. RNA cleavage assays

RNA cleavage assays were performed according to a previously described procedure [38].

#### 2.1.3. Gel imaging and analysis

The data were collected using an Amersham™ Typhoon™ (Cytiva, Washington, D.C., USA) and quantified using MultiGauge 3.0 software (Fujifilm, Minato, Tokyo, Japan). In the case of all diagrams, error bars represent SD values calculated based on three independent experiments.

#### 2.1.4. Tertiary structure prediction

The tertiary structure of the hDicer helicase K70A mutant was obtained using SWISS-MODEL (http://swissmodel.expasy.org) [41]. Tertiary structures of pre-miRNAs were predicted using the RNA Composer server (https://rnacomposer.cs.put.poznan.pl) [42].

#### 2.1.5. SAXS

Small Angle X-ray Scattering (SAXS) measurements were performed at the P12 beamline of the PETRA III storage ring at the DESY (Deutsches Electron Synchrotron) in Hamburg, Germany. SEC-SAXS technique was used to increase the sample quality. The P12 beamline was equipped with an Agilent 1260 Infinity II Bio-inert liquid chromatography system (LC Agilent, Waldbronn, Germany). The data were recorded as a sequential set of 2880 individual 1 s frames corresponding to one column volume for each protein sample (48 min total). Each individual 2D image underwent data reduction (azimuthal averaging) and normalization to the intensity of the transmitted beam to generate 1D scattering profiles plotted as *I(s) vs s* through the momentum transfer range of 0.05<s<6 nm-1(where s=4πsinθ/λ and 2θ is the scattering angle). The program CHROMIX was integrated into the automated data processing pipeline. Automated CHROMIXS selection of frames recorded before or after the sample peak, corresponding to the buffer, were averaged and used for the subtraction of the background scattering contribution from the sample frames [43]. The ATSAS package was used for further data analysis and modelling [44]. The quaternary structures of pre-miRNA•HEL complexes were modeled using SASREF modelling software [45]. The spatial structures of pre-miRNAs were predicted using RNAcomposer. Theoretical calculations of the structural parameters (radius of gyration, volume) were performed in CRYSOL.

## 3. Results

### 3.1. The helicase domain of hDicer is responsible for the ATP hydrolysis activity of hDicer

To investigate the biochemical properties of the hDicer helicase domain, we expressed this domain (called HEL) using *Escherichia coli* (Supplementary **Fig. S1 a**). Because the hDicer helicase domain contains the well-conserved Walker A and Walker B motifs (**Fig. 1c**), which are associated with ATP binding and hydrolysis, respectively, in the first step, we tested whether the obtained HEL preparation can hydrolyze ATP. We examined a time-course of ATP hydrolysis using equimolar amounts of HEL and [γ^32^P]-ATP substrate (2 nM). The reaction mixtures were separated by denaturing polyacrylamide gel electrophoresis (PAGE) and visualized by phosphorimaging. We observed that under the applied reaction conditions, HEL consumed half of the substrate in ∼40 minutes (**Fig. 2a, 2d**). Since ATPase activity has not previously been associated with hDicer, we confirmed this result using ATP hydrolysis assays involving full-length wild-type hDicer (WT hDicer) (Supplementary **Fig. S1 b**). We compared activity to a hDicer mutant with an lysine to alanine substitution in the Walker A motif at position 70 (hDicer_K70A) (Supplementary **Fig. S1 c**) and a hDicer variant lacking the helicase domain (hDicer_ΔHEL) (Supplementary **Fig. S1 d**). The well-conserved lysine (K) residue in the Walker A motif is crucial for ATP binding, and mutations of this residue in all investigated ATPases/helicases strongly inhibited nucleotide binding and enzymatic activity [46]. Our data revealed that ATP was hydrolyzed in reactions carried out with WT hDicer (**Fig. 2b**), and that the efficiency of this hydrolysis was similar to the efficiency of ATP hydrolysis by HEL (**Fig. 2d**). Very low levels of ATP hydrolysis activity were observed for hDicer_K70A (**Fig. 2c**), while no ATP hydrolysis was detected for hDicer_ΔHEL (Supplementary **Fig. S2**). A comparative analysis of ATP hydrolysis activity between WT hDicer and hDicer_K70A revealed that hDicer_K70A hydrolyzed ATP with ∼400-times lower efficiency than WT hDicer (Supplementary **Fig. S3**).

**Fig. 2.**
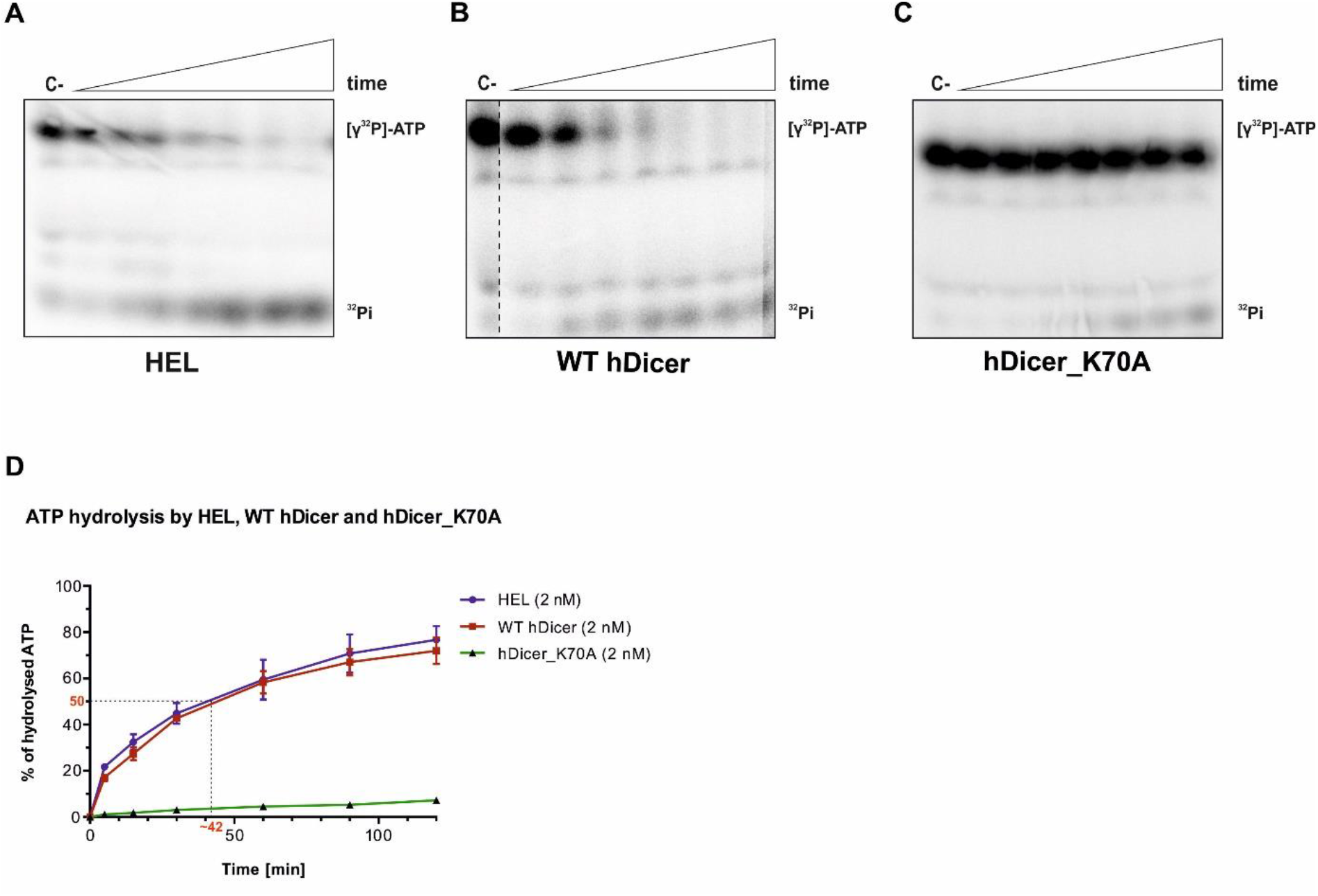
Time-dependent ATP hydrolysis activity of HEL, WT hDicer and hDicer_K70A. **a** Denaturing PAGE analysis of mixtures of [γ^32^P]-ATP (2 nM) and HEL (2 nM). Samples were taken at: 1, 5, 15, 30, 60, 90 and 120 min from reaction mixtures incubated at 37 °C, C-indicates a control sample with no protein and ^32^Pi indicates the product of [γ^32^P]-ATP hydrolysis. **b** Denaturing PAGE analysis of mixtures of [γ^32^P]-ATP and WT hDicer (2 nM). Samples were analyzed at: 1, 5, 15, 30, 60, 90 and 120 min, reaction mixtures were incubated at 37 °C, C-indicates a control sample with no protein and ^32^Pi indicates the product of [γ^32^P]-ATP hydrolysis. **c** Denaturing PAGE analysis of the mixtures of [γ^32^P]-ATP and hDicer_K70A (2 nM). Samples were taken at: 1, 5, 15, 30, 60, 90 and 120 min form reaction mixtures incubated at 37 °C, C-indicate a control sample with no protein and ^32^Pi indicate the product of [γ^32^P]-ATP hydrolysis. **d** Graph quantifying progression of ATP hydrolysis reactions. The *x*-axis represents the incubation time expressed in minutes, and the *y*-axis represents the percentage of hydrolyzed ATP by HEL, WT hDicer and hDicer_K70A. Error bars represent standard deviations (SD) based on three separate experiments.

Previous research has shown that K70A mutation in hDicer does not affect its RNA cleavage activity [46]. To ensure that the observed differences in ATP hydrolysis activity did not result from differences in the quality or folding of recombinant proteins, we assessed the RNase activity of WT hDicer, hDicer_K70A and hDicer_ΔHEL. The time-course cleavage assay involved 5′-^32^P-labeled pre-miRNA (5 nM) and the protein (10 nM). The results showed that, under the applied reaction conditions, WT hDicer, hDicer_K70A and hDicer_ΔHEL displayed similar pre-miRNA cleavage efficiency (Supplementary **Fig. S4**). Overall, the results indicate that the observed differences in ATP hydrolysis activity between WT hDicer, hDicer_K70A and hDicer_ΔHEL did not result from the quality of the proteins used in the assay.

Based on comparing the tertiary structures of the helicase domain of hDicer and the PcrA helicase from *Geobacillus stearothermophilus* (**Fig. 3a**), it can be inferred that lysine 70 in the Walker A motif of hDicer might be involved in ATP binding (**Fig. 3b**). Thus, we hypothesized that the diminished ATP hydrolysis activity of hDicer_K70A might be associated with reduced ATP-binding. To test this hypothesis we performed ATP binding assays involving either WT hDicer or hDicer_K70A. We performed an electrophoretic mobility shift assay (EMSA) with reaction mixtures containing: [γ^32^P]-ATP substrate (2 nM) and WT hDicer (3.59, 7.19, 14.38, 28.75, 57.5, 115 nM) or hDicer_K70A (36, 72, 144, 288, 575, 1150 nM). The data showed that WT hDicer bound ATP with a K_d_ value of ∼85 nM, while only weak ATP binding could be observed for hDicer_K70A. This weak binding precluded determining a K_d_ value for this hDicer variant (Supplementary **Fig. S5**). Accordingly, our data suggest that the lower level of ATP hydrolysis activity of hDicer_K70A results from its reduced affinity for ATP.

**Fig. 3.**
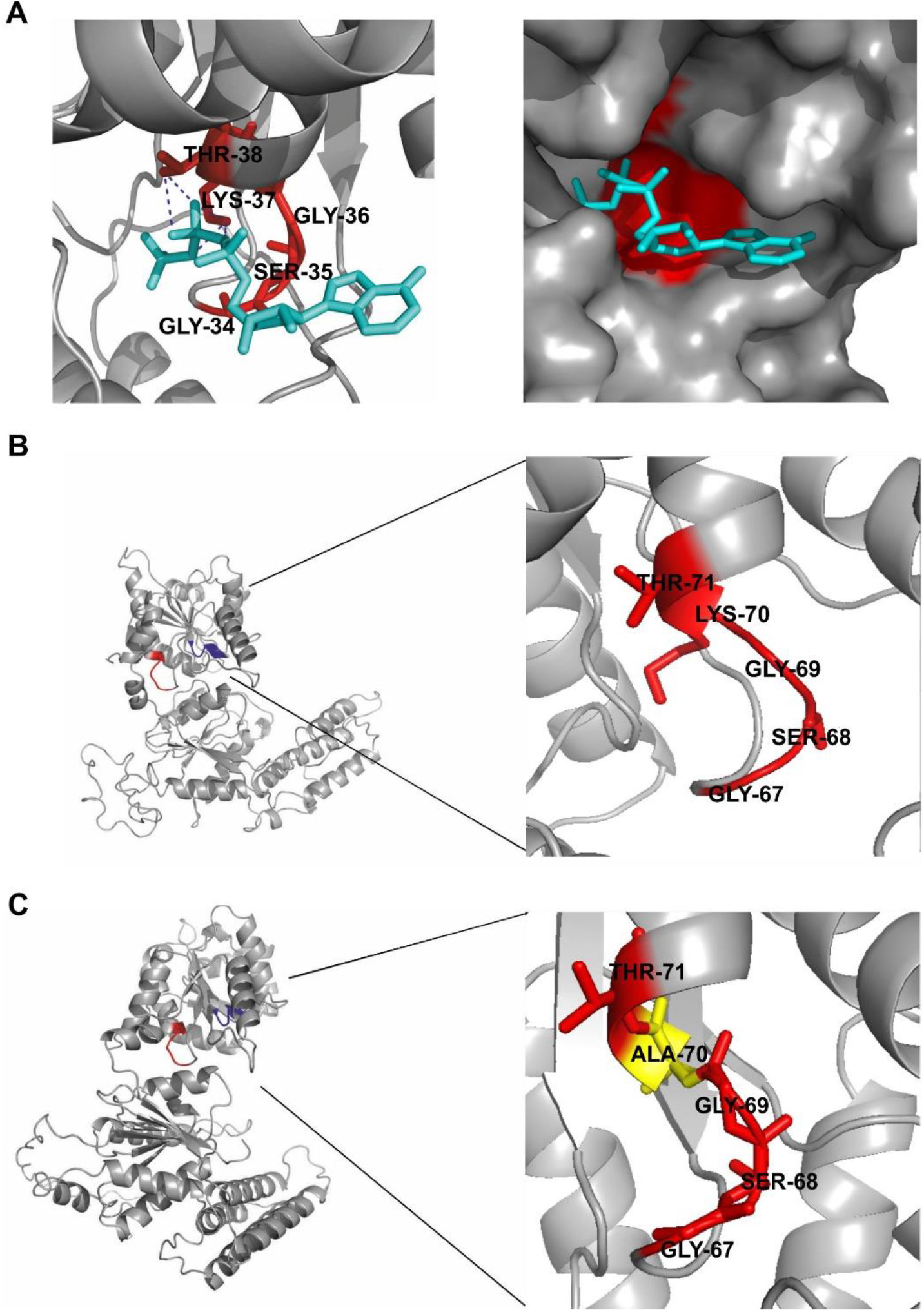
Walker A motif structures in hDicer and related proteins. **a** Structure of the Walker A motif of PcrA helicase from *Geobacillus stearothermophilus* with interacting ATP substrate. The image was generated in PyMOL, based on structural data for the PcrA helicase (PDB entry 3PJR) [66]; the Walker A motif (GSGKT) is marked in red and the ATP is marked in turquoise. Hydrogen bonds between threonine 38 and lysine 37 and phosphate are indicated. Structure of the Walker A motif of: **b** WT hDicer helicase domain and **c** hDicer_K70A helicase domain. Tertiary structures of the WT hDicer helicase and hDicer_K70A variant helicase domain were obtained using SWISS-MODEL based on the structural data for hDicer (PDB entry 5ZAL) [9]. The fragment encompassing amino acid residues 67-71 of the Walker A motif (GSGKT) is marked in red. In the hDicer_K70A variant, the respective region (GSGAT) is indicated.

Importantly, all presented studies were carried out under low-turnover conditions; thus, as a next step we analyzed the ATP hydrolysis activity of WT hDicer and HEL under high-turnover conditions. In this ATP hydrolysis assay we used 1 μM ATP substrate and 2 nM WT hDicer or HEL. However, under the applied reaction conditions, we did not observe the products of ATP hydrolysis (Supplementary **Fig. S6**).

These results show that hDicer can hydrolyze ATP and that the domain responsible for this activity is the helicase domain. To the best of our knowledge, this is the first time this activity has been observed. The ATPase activity of hDicer can be observed under low-turnover conditions.

### 3.2. Nucleic acid binding activity of the hDicer helicase domain

Previous work has revealed that Dicer enzymes can bind several RNA substrates in the cell, and that binding of some of RNAs occurs without cleavage [35]. The authors have suggested that such passive binding of substrates is mediated by the Dicer helicase domain [35]. Assuming that the hDicer helicase domain plays an important role in the binding of various hDicer substrates, we investigated the nucleic acid binding properties of this domain. The binding assays contained the following types of substrates: (i) single-stranded RNAs (ssRNAs), including R10 (12 nt), R20 (21 nt), R30 (32 nt), R40 (42 nt) and R50 (56 nt); (ii) corresponding single-stranded DNAs (ssDNAs): D10, D20, D30, D40 and D50; (iii) pre-miRNAs, including pre-mir-21, pre-mir-33a and premir-16-1; (iv) dsRNAs, including dsRNA_blunt (32 bp) and dsRNA_over (30 bp with 2-nt 3′-overhangs); and (v) corresponding dsDNAs: dsDNA_blunt and dsDNA_over. Single-stranded substrates were ^32^P-labeled at the 5′ end and double-stranded substrates contained one strand that was 5′-^32^P-labeled. Before they were applied to the reaction mixtures, ssRNAs and ssDNAs were denatured at 95 °C for 3 min and placed on ice to ensure they were single-stranded. Reaction mixtures containing a substrate (∼2.5 nM) and HEL dilutions (2.97, 5.94, 11.86, 23.75, 47.5, 95 µM) were incubated at room temperature for 15 min. They were then separated using EMSA and visualized by phosphorimaging (**Fig. 4**). In general, the observed band patterns indicated that HEL can bind 20nt ssRNAs and 20-nt ssDNAs, as well as longer substrates (**Fig. 4**). However, in the case of ssRNAs (**Fig. 4a**), distinct but low abundant complexes were only observed for R20 and R30 molecules. RNA•HEL complexes with R40 and R50 molecules were poorly detected. The ambiguous results and the weak signal made it impossible to determine the K_d_ values for the ssRNA•HEL complexes. In the case of ssDNAs (**Fig. 4b**), we observed distinct ssDNA•HEL complexes for the D30, D40 and D50 substrates. The densitometry analyses allowed us to calculate K_d_ values for the D40•HEL and the D50•HEL complexes; these values were: ∼8.5 µM for D40•HEL, and ∼7.4 µM for D50•HEL.

**Fig. 4.**
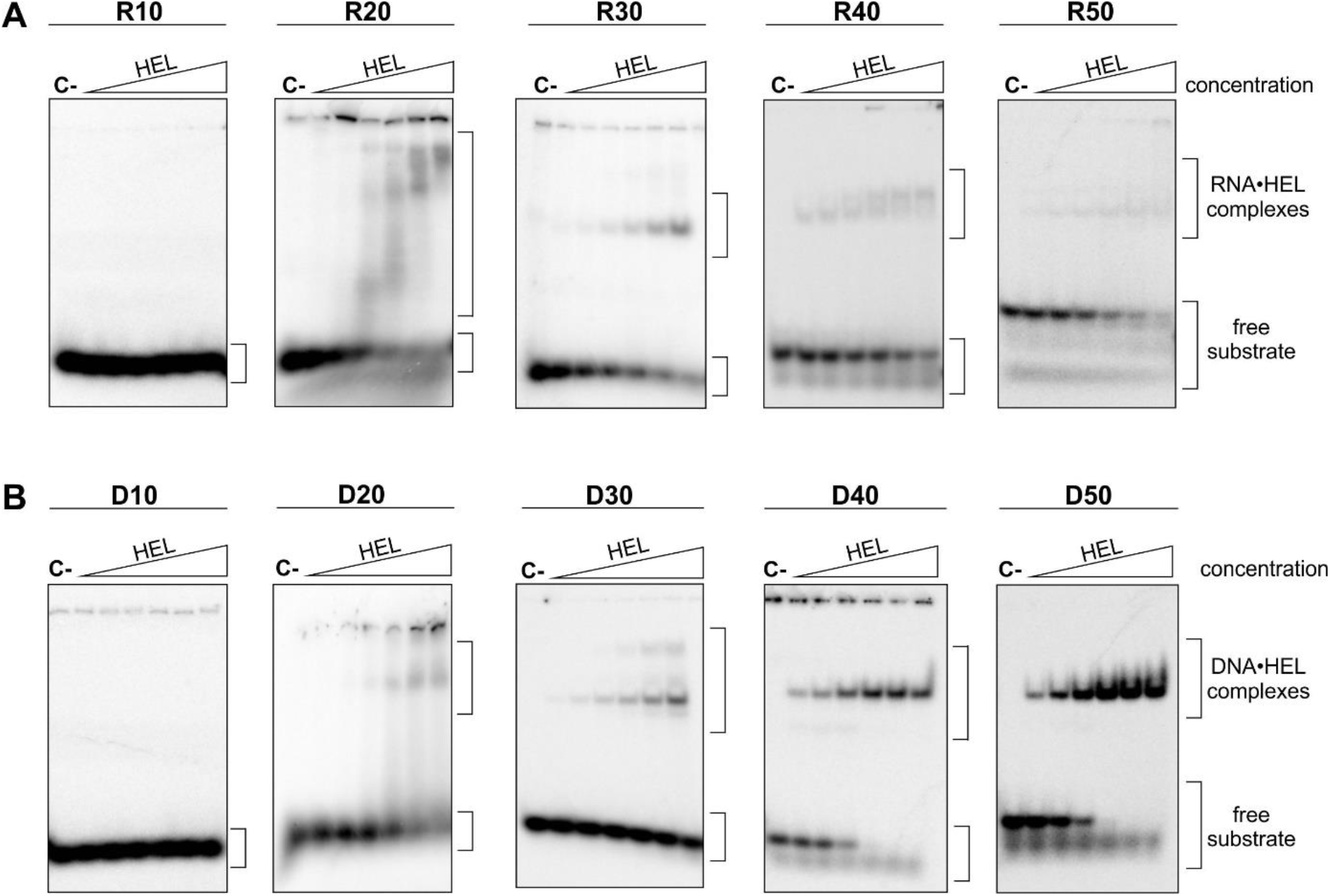
Nucleic acid binding activity of HEL. **a** EMSA with HEL and 5′-^32^P-labeled single-stranded RNAs (ssRNAs) (2.5 nM): R10, R20, R30, R40, R50. **b** EMSA with HEL and 5′-^32^P-labeled single-stranded DNAs (ssDNAs) (2.5 nM): D10, D20, D30, D40, D50. Increasing amounts of HEL (2.97, 5.94, 11.86, 23.75, 47.5, 95 µM) are represented by a triangle. Reaction mixtures were incubated at room temperature for 15 min. C-indicates a control sample with no protein.

Taking into consideration the possibility that binding between a nucleic acid and HEL might be unstable during EMSA analysis, we next applied bio-layer interferometry (BLI) to investigate the interactions between ssRNAs or ssDNAs and HEL. BLI is an optical method for studying the affinity between molecules in real time without the need for fluorescence or radioisotope-labeled particles. This method is based on using biosensors that are specific to the proteins or other substrates being tested. For example, biosensors with Ni-NTA beads are suitable for use with proteins that have His6-tags, as in the case of HEL. BLI can be used to measure association and dissociation rate constants, as well as the equilibrium dissociation constant (K_d_). Given the lack of observed complexes for R10•HEL and D10•HEL (**Fig. 4**) in the BLI assay, we used the following ssRNAs: R20, R30, R40, R50. Additionally, we used the following ssDNAs: D20, D30, D40, D50. The measurements were carried out using HEL (1 µM) and several substrate dilutions (3.125, 6.25, 12.5, 25, 50 and 100 µM). First, HEL was immobilized on Ni-NTA biosensor. After that, HEL-loaded sensor was immersed in a ligand-containing well, to monitor the association, and in a buffer-containing well, to monitor the dissociation of the nucleic acid•protein complex. Each measurement was repeated three times. The signals were too low to calculate reliable K_d_ values for the following substrates: R20, R30, D20 and D30 (Supplementary **Fig. S7**), although the association and dissociation curves were recorded. K_d_ values that met the quality control criteria, calculated based on association and dissociation curves, were collected for: R50•HEL (∼23 µM) (Supplementary **Fig. S7 a**), D40•HEL (∼22 µM) and D50•HEL (∼21 µM) (Supplementary **Fig. S7 b**). Observed differences in K_d_ values obtained using EMSA and BLI methods may result from differences between the two different approaches. For example, the BLI method is more sensitive, allowing association and dissociation of complexes to be monitored in real time. Moreover, in the BLI method we applied increasing amounts of substrate, while in EMSA we used increasing amounts of protein. Nevertheless, it is important to note that the K_d_ values estimated by both methods were in a similar micromolar range.

The next set of binding assays involved pre-miRNA substrates: pre-mir-21, pre-mir-33a and pre-mir-16-1. The selected pre-miRNAs differ in the compactness of their secondary structures: pre-mir-21 adopts a compact structure, with a small terminal loop, pre-mir-33a contains large internal loops and bulges, but has a small terminal loop, while pre-mir-16-1 has a more relaxed structure, with a 9-nt apical loop (**Fig. 5a**). Reaction mixtures containing a 5′-^32^P-labeled substrate (∼2.5 nM) and different HEL dilutions (11.86, 23.75, 47.5, 95 µM) were incubated at room temperature for 15 min. Mixtures were then separated using EMSA and visualized by phosphorimaging. The results of the EMSA experiment revealed smeared bands for all reaction sets (**Fig. 5b**). Band smearing can be attributed to weak and unstable binding between the pre-miRNA and HEL. The obtained results did not allow calculation of K_d_ values for the pre-miRNA•HEL complexes.

**Fig. 5.**
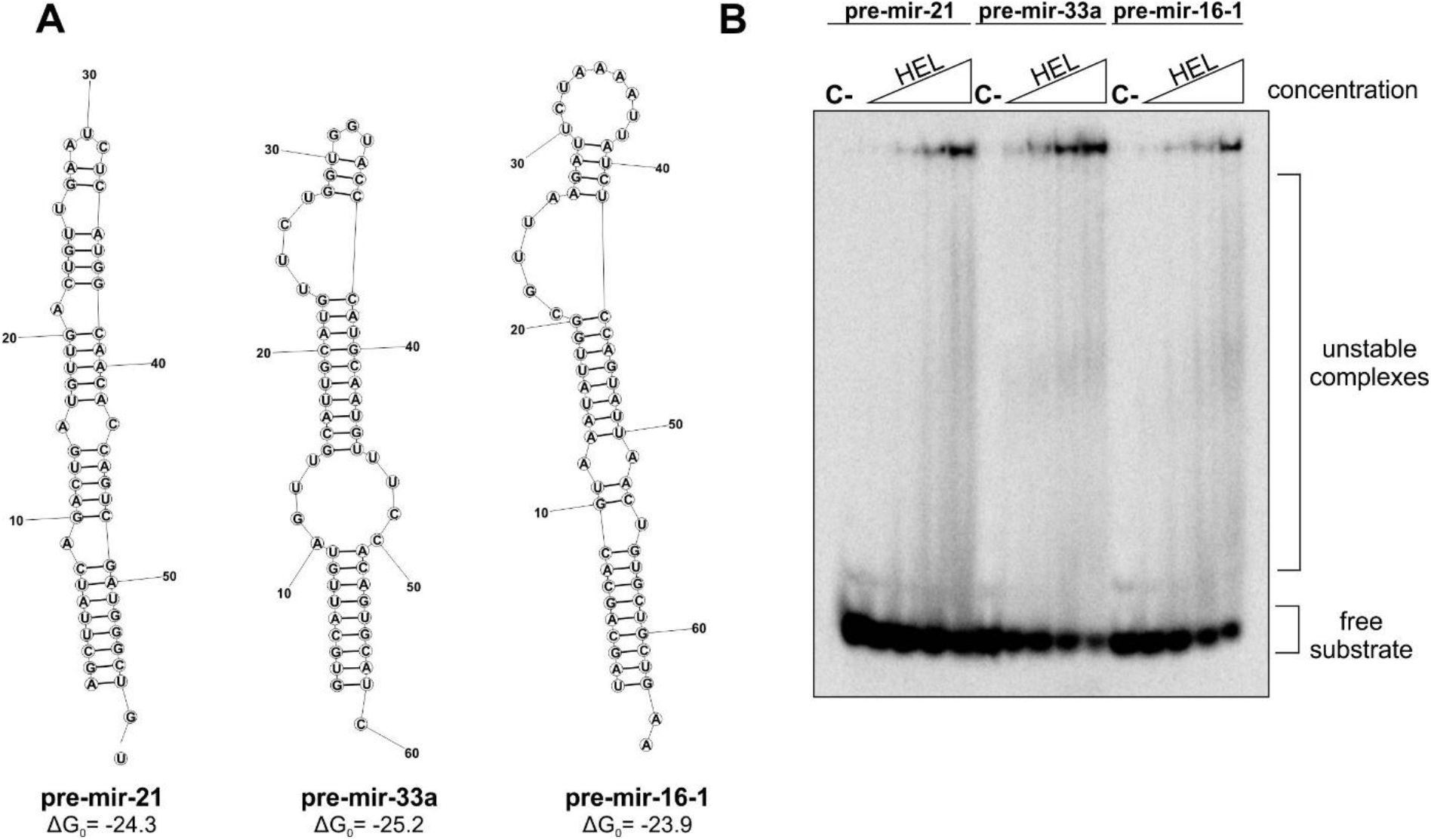
Pre-miRNA binding by HEL. **a** Secondary structures of pre-miRNAs used in the study. The predicted structures for pre-mir-21, pre-mir-33a, and pre-mir-16-1 generated using the RNAstructure Fold online tool (Mathews Lab) [47]. The structures are color-annotated according to base-pairing probability. Free energy values expressed in kcal/mol are shown at the bottom. Nucleotides are numbered starting from the 5’-end. **b** EMSA with HEL and 5′-^32^P-labeled pre-miRNAs (2.5 nM): pre-mir-21, pre-mir-33a, pre-mir-16-1. Increasing amounts of HEL (11.86, 23.75, 47.5, 95 µM) are represented by a triangle. Reaction mixtures were incubated at room temperature for 15 min. C-indicates a control sample with no protein.

To get better insight into possible interactions between pre-miRNAs and HEL we applied small-angle X-ray scattering (SAXS) analysis. In this analysis we used pre-mir-16-1 and pre-mir-21. We used SASREF software [45] to model structures of pre-miRNA•HEL complexes using our experimental SAXS curves (Supplementary Datasets related to the SAXS studies), known structural data for HEL (PDB entry 5ZAL), and structures of pre-mir-16-1 and pre-mir-21 (predicted by RNAComposer) (**Fig. 6**; the corresponding structural data are in Supplementary **Table S1**). Analysis of structural parameters such as volume and molecular weight revealed that pre-miRNA•HEL complexes formed with a 1:1 ratio. The obtained models indicated that HEL contacts the apical regions of pre-miRNAs mostly through the HEL2i subdomain. It must be underlined, however, that the above-mentioned assays, presented in **Fig. 5** and **Fig. 6**, involved a stand-alone helicase domain. The helicase domain in the context of full-length hDicer might operate differently.

**Fig. 6.**
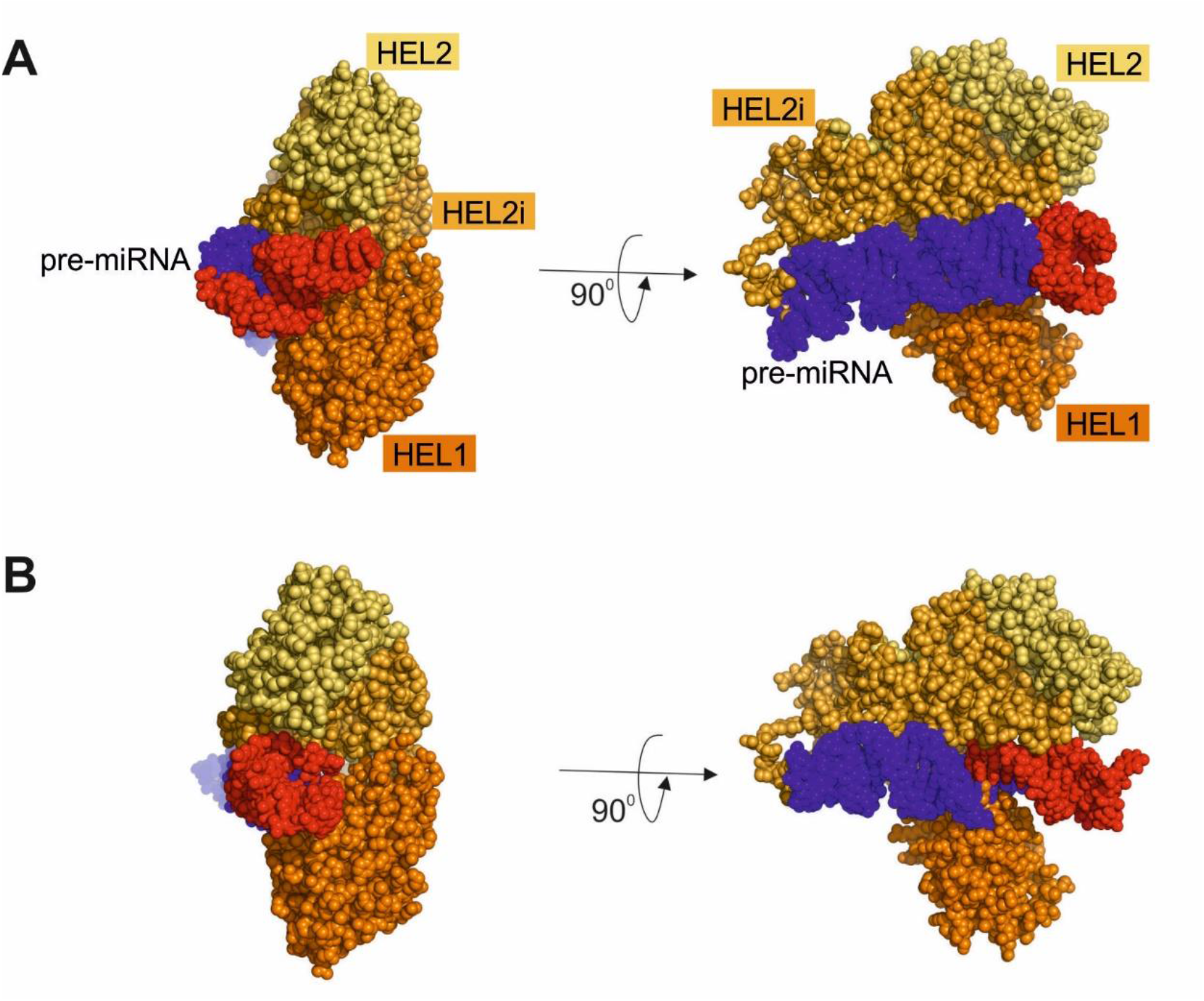
SASREF rigid body models of the hDicer helicase domain complexes with **a** pre-mir-16-1 and **b** pre-mir-21. Structure of the hDicer helicase domain (PDB entry 5ZAL) was generated using SWISS MODEL and structures of pre-miRNAs were predicted using RNAComposer. The three helicase subdomains are distinguished: HEL1 (bright orange), HEL2i (light orange), HEL2 (yellow). In the pre-miRNA structures, stem regions are indicated in purple, while the apical loop regions (nucleic acid residues 20-40) are indicated in red.

Finally, binding assays involving dsRNA and dsDNA substrates were carried out. We analyzed binding to a 32-bp RNA or DNA duplex (blunt) and a 30-bp RNA or DNA duplex with a 2-nt 3′ overhang on each end (over). Thus, in total, we tested four types of substrates: dsRNA_blunt, dsRNA_over, dsDNA_blunt, and dsDNA_over. Double-stranded substrates (∼2.5 nM) were incubated with HEL dilutions (2.97, 5.94, 11.86, 23.75, 47.5, 95 µM) at room temperature for 15 min. EMSA revealed no band-shifts for the tested sets; this indicated that HEL does not bind double-stranded nucleic acids (Supplementary **Fig. S8**).

Altogether, the results reveal that HEL can bind single-stranded RNAs and DNAs of 20-nt and longer but it does not interact with dsRNA and dsDNA substrates. We also found that pre-miRNA substrates, which inherently contain partially double-stranded regions, do not form stable complexes with the stand-alone helicase domain of hDicer.

### 3.3. The hDicer helicase domain rearranges the structure of interacting RNAs in an ATP-independent manner

The results of the EMSA assay revealed weak binding between the ssRNA substrates (R20, R30, R40 and R50) and HEL. Moreover, these results also showed a HEL-concentration-dependent loss of the main substrate form (the most intense band, corresponding to the dominant substrate form) (**Fig. 4a**). With this in mind, we hypothesized that HEL might rearrange the structures of the interacting ssRNA substrates or trigger their degradation. To test these hypotheses, we analyzed the substrates and reaction products of the binding reactions by (i) comparative PAGE, under native and denaturing conditions, and (ii) circular dichroism spectroscopy (CD). All experiments were carried out using the R40 substrate and HEL.

As in the binding assays above, reactions contained ^32^P-labeled R40 (∼2.5 nM) and different HEL dilutions (2.97, 5.94, 11.86, 23.75, 47.5, 95 µM). Native PAGE confirmed inefficient formation of complexes between R40 and HEL (**Fig. 7a**). We also confirmed the existence of HEL-concentration-dependent loss of the main substrate conformation. More specifically, there was a reduction in the slow-migrating conformer of R40, presumably representing a single-stranded form of R40. This change was either accompanied by an increase in quantity of the fast migrating conformers of R40 (presumably representing more compact conformers of R40) or products of R40 degradation.

**Fig. 7.**
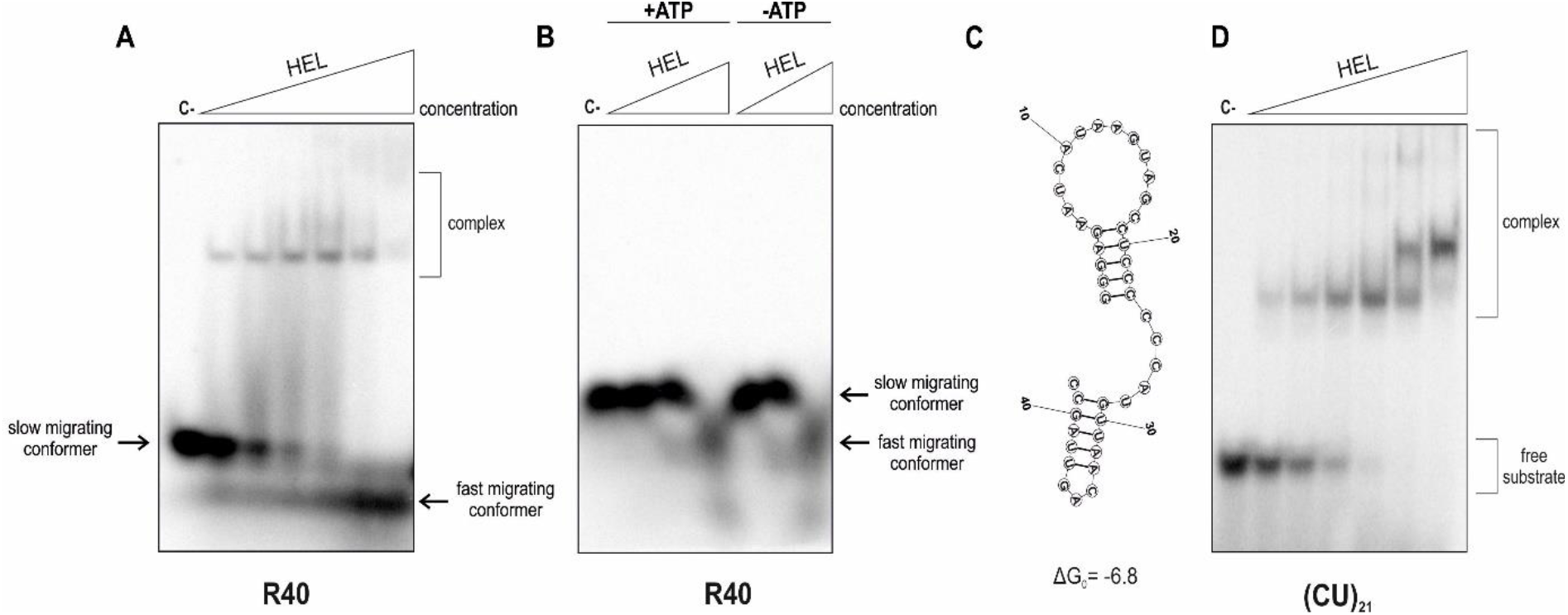
Interactions between 42-nt RNAs and HEL. **a** EMSA with HEL and 5′-^32^P-labeled R40 (42-nt) (2.5 nM). Increasing amounts of HEL (2.97, 5.94, 11.86, 23.75, 47.5, 95 µM) are represented by a triangle. Reaction mixtures were incubated at room temperature for 15 min. C-indicates a control sample with no protein. **b** Native PAGE analysis of mixtures of 5′-^32^P-labeled R40 (2.5 nM) and increasing amounts of HEL (5.94, 23.75, 95 µM). After a 15 min incubation at room temperature, sodium dodecyl sulfate (SDS) to a final concentration of 1% was added to denature protein. C-indicates a control sample with no protein. +ATP indicates reaction mixtures with 1 mM ATP. -ATP indicates reaction mixtures without ATP. **c** Secondary structure of R40 generated using the RNAstructure Fold online tool (Mathews Lab) [47]. The structure is color-annotated according to base pairing probability. The free energy value expressed in kcal/mol is shown at the bottom. Nucleotides are numbered starting from the 5’-end. **d** EMSA with HEL and 5′-^32^P-labeled (CU)_21_ (42-nt) (2.5 nM). Increasing amounts of HEL (2.97, 5.94, 11.86, 23.75, 47.5, 95 µM) are represented by a triangle. Reaction mixtures were incubated at room temperature for 15 min. (C-) − a control sample with no protein.

To test the integrity of the substrate after interaction with HEL, we separated the reaction mixtures, including R40 and HEL, under denaturing conditions (Supplementary **Fig. S9**). Our analysis revealed that R40 stayed intact even when the highest concentration of HEL was applied (95 µM). These results argue that R40 was not degraded when incubated with HEL. In the next binding assay we tested the influence of ATP on changes in R40. In addition, after 15 min incubation of R40 with HEL, with or without ATP, SDS was added to the reaction mixtures, to final concentration of 1%. SDS disrupts interactions between proteins and nucleic acids by denaturing the protein but leaving the nucleic acid structurally intact [32]. Native PAGE showed that under the applied reaction conditions with increasing HEL concentrations, the slower-migrating conformer of R40 gradually disappeared while faster-migrating conformers of R40 increased in intensity (**Fig. 7b**). The output of the assay did not depend on ATP.

Altogether, the collected results indicated that HEL influences R40 structure, and that this process is ATP-independent. We hypothesized that upon HEL binding, R40 can adopt a more compact structure. A possible secondary structure of R40 was predicted using the RNA structure web server [47] (**Fig. 7c**). We noticed that the secondary structure of R40 contains double-stranded regions and, as indicated in **Fig. 5**, substrates with double-stranded regions, such as pre-miRNAs, are poorly bound by HEL. To further test the hypothesis that HEL can rearrange the structure of interacting RNAs, we performed the HEL binding assay using a substrate that cannot adopt a secondary structure: a 42-mer composed of (CU)_21_ repeats. This time, with increased HEL concentration, we noticed clear distinct bands correlating to RNA•HEL complexes, and no appearance of the fast-migrating conformers (**Fig. 7d**). These results further proved that HEL can rearrange the structure of interacting RNAs, providing that RNA has the intrinsic potential to adopt secondary structures. Further examples of HEL-assisted rearrangements of interacting RNA structure can be found in Supplementary **Fig. S10**.

Next, we used CD spectroscopy for further analysis of the effect of HEL on the R40 structure. CD spectroscopy is a simple optical technique that is most sensitive to the structural polymorphism of nucleic acids and proteins [48]. Since characteristic bands for nucleic acids and proteins are separated in the CD spectrum [49], an independent structural analysis of nucleic acids and proteins can be performed. Analyzing the shape of a CD spectrum provides information on the structure of a biomolecule [50] [51]. In the experiment, R40 (28 µM) was incubated in a buffer solution (10 mM HEPES pH 8.0, 300 mM NaF) with or without HEL protein (28 µM). Alternatively, HEL (28 µM) was incubated in the buffer solution alone. A comparison of the shapes of the CD spectra, in a spectral range of 210 to 350 nm, is presented in **Fig. 8a**. The helicase domain of hDicer is mostly composed of alpha helices (**Fig. 1c**), secondary structures which typically give a negative peak at 222 nm [50]. The double-stranded helical regions of RNA give a positive peak at ∼270 nm [50]. Our results confirmed that the CD spectrum of the HEL protein had a negative minimum value at 222 nm. In contrast, the CD spectrum of R40 alone (**Fig. 8a**) had a maximum value at ∼270 nm, which is characteristic for dsRNA structures that can be found in R40 (**Fig. 8c**) [49]. The CD spectrum of the R40•HEL complex had a negative minimum value typical for a protein with a dominant alpha helical structure (at 222 nm); however, the maximum value for dsRNA, at ∼270 nm, was flattened (**Fig. 8a**). This result may be due to interactions between HEL and R40 and structural rearrangements of R40 upon HEL binding, such as loss of double-stranded helical regions in R40.

**Fig. 8.**
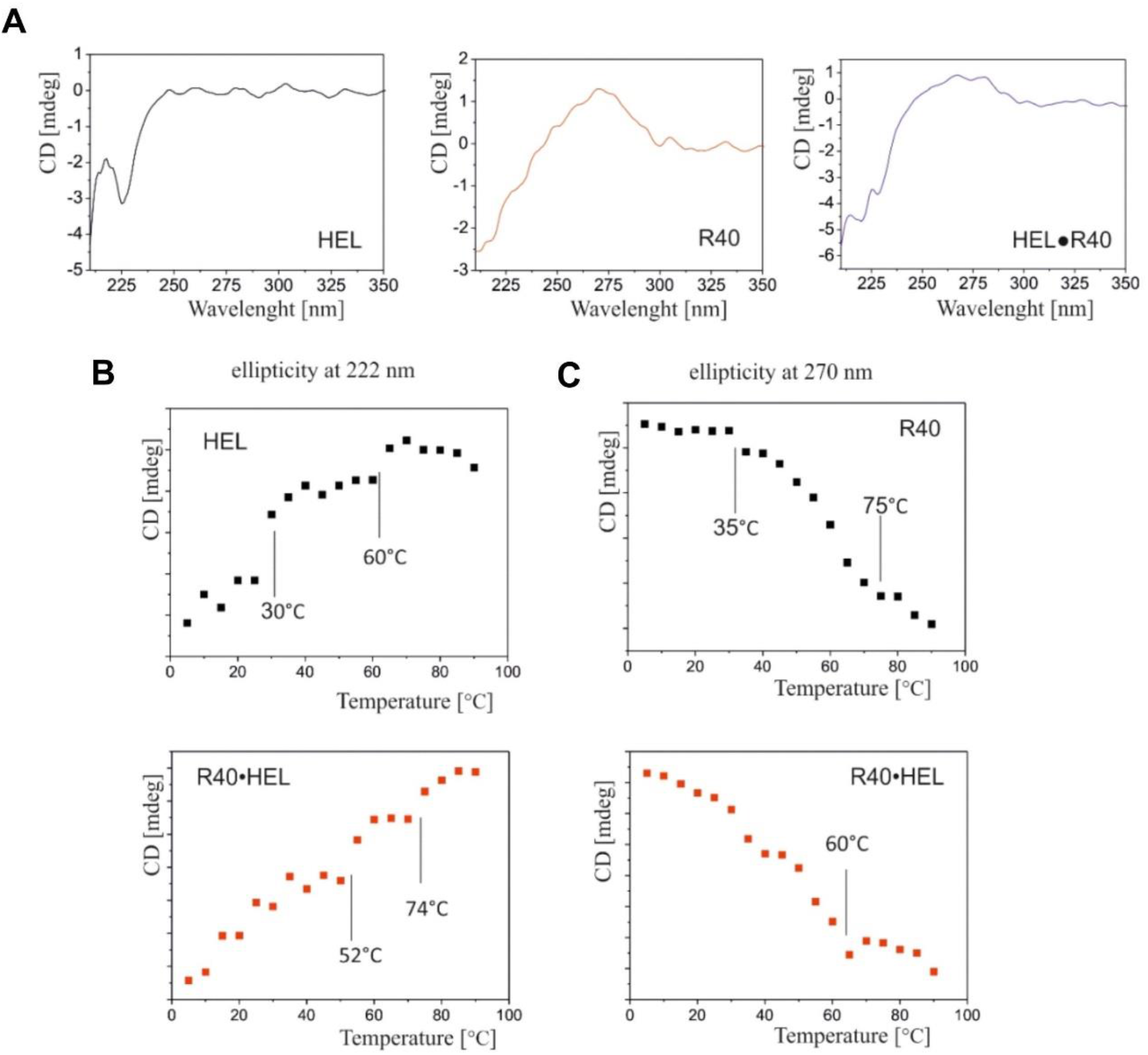
Circular dichroism (CD) analysis of R40, HEL, and the HEL•R40 complex. **a** Example CD spectra for: HEL (black), R40 (red), and the HEL•R40 complex (blue). Measurements were carried out at 15 °C. **b** CD thermal analysis at 222 nm for HEL (black) and the HEL•R40 complex (red). **c** CD thermal analysis at 270 nm for R40 (black) and the HEL•R40 complex (red).

Next, a thermal analysis was carried out to analyze the alpha-helical content in the HEL and R40•HEL samples (at 222 nm), and the dsRNA content in the R40 and R40•HEL samples (at 270 nm). For the HEL sample (**Fig. 8b**), thermal analysis was carried out at 222 nm. As the temperature increased (from 5 to 90 °C), an increase in peak intensity was observed. Moreover, two discontinuities were visible at ∼30 °C and ∼60 °C. These discontinuities are called “phase–transition temperatures” [52], and are associated with the structural changes that occur in a protein as the temperature increases. In other words, as the temperature increases, the secondary structure content of a protein decreases. For the R40•HEL complex (**Fig. 8b**, red dotted curve), the observed discontinuities were shifted towards higher temperatures (i.e., at 52 °C and 74 °C, respectively). This indicates that the structure of HEL stabilized after the R40 binding occurred.

To investigate the structural changes in R40, thermal analysis was carried out at 270 nm (**Fig. 8c**). In the case of the R40 sample, as the temperature increased, a decrease in peak intensity was observed, which was related to the melting of double-stranded structures within R40. Moreover, for the R40 sample (**Fig. 8c**, black dotted curve), two discontinuities were observed at 35 °C and 75 °C. For the R40•HEL complex (**Fig. 8c**, red dotted curve), the discontinuity at ∼35 °C was barely visible, and the discontinuity observed at 75 °C, for R40 alone, shifted towards a lower temperature, 60 °C. Thus, the difference in phase transition temperatures between R40 alone and R40 complexed with HEL was 15 °C. A comparison of the experimental and theoretical thermodynamic data, examining to R40 and R40•HEL samples, can be found in Supplementary **Table S2**. The thermodynamic parameters reveal that the entropy and enthalpy values for the R40•HEL complex decreased by ∼12% compared to the corresponding values for R40 alone. Collectively, these results imply that the structure of R40 shifted toward a more relaxed state (lower dsRNA content) upon HEL binding.

In summary, the collected data support the hypothesis that the helicase domain of hDicer can induce conformational changes in the RNAs with which it interacts.

## 4. Discussion

The helicase domain of hDicer has well-conserved ATPase motifs; however, so far, there have been no studies demonstrating the ATPase activity of hDicer. The data in this manuscript show for the first time that hDicer is capable of ATP hydrolysis (**Fig. 2**). Recently published data suggest that ancestral Dicers in the hDicer lineage had decreased ATP binding affinity and thus lost ATPase activity, in contrast to plant or arthropod Dicers [23]. Our study shows that hDicer can bind ATP with a K_d_ of ∼85 nM (Supplementary **Fig. S5**). Mutation of the well-conserved lysine (K70) to alanine in the ATP-binding motif (the Walker A motif) of the hDicer helicase domain drastically reduced ATP binding of the hDicer_K70A variant (Supplementary **Fig. S5**), resulting in a ∼400-fold decrease in ATPase activity of this hDicer variant (Supplementary **Fig. S3**). It is important to underline that in a previous study [23], ATP hydrolysis assays were carried out under the high-turnover conditions, with a high excess of a substrate over the enzyme (500:1 molar ratio). In our assays, we used approximately equimolar amounts of the substrate and enzyme. When we used a 1:500 molar ratio of enzyme to substrate, we did not detect products of ATP hydrolysis in reactions with the hDicer helicase domain or with full-length hDicer (Supplementary **Fig. S6**). It is possible that under excess substrate conditions, the products of ATP hydrolysis might inhibit the ATPase activity of the helicase domain [53, 54]. Moreover, published data reveal that RNA may stimulate ATPase activity of ecdysozoan invertebrate Dicers by increasing their ATP affinity [23]. Our study shows that RNA does not influence ATP binding by the wild-type hDicer (Supplementary **Fig. S11**). To summarize, preservation of ATPase activity by hDicer suggests that this enzyme might be involved in currently unknown cellular pathways that utilize this biochemical function. We may speculate that activation of the hDicer ATP-hydrolyzing function may trigger conformational changes of hDicer’s domains. Then, a stimulated hDicer may transmit downstream signals, resulting in the activation or repression of yet undefined cellular factors. In this way hDicer might be directly involved in signal transduction.

Considering hDicer interactions with its canonical substrates, pre-miRNAs, the ends of the pre-miRNA substrate are bound by the PPC cassette of hDicer [7] while the single-stranded apical loop region of pre-miRNA interacts with the helicase domain [9]. However, in the cell, hDicer may also interact with RNAs other than pre-miRNAs, e.g., with mRNAs or lncRNAs [35]. Binding of these types of RNA is likely initiated by the helicase domain of hDicer [35]. These observations prompted us to investigate the substrate specificity of the hDicer helicase domain. The *in vitro* binding assays revealed that the helicase domain was capable of binding ∼20-nt ssRNAs, ssDNAs, and longer substrates (**Fig. 4**). However, the binding of pre-miRNAs, which are partially double-stranded, was very inefficient (**Fig. 5**), as was the binding of dsRNA and dsDNA by the hDicer helicase (Supplementary **Fig. S8**). The most likely explanation for these observations are the structural constraints of the hDicer helicase domain [9]. A comparison of the tertiary structures of *A. thaliana* DCL1 [55], *D. melanogaster* Dicer-2 [56], and hDicer [9] indicates that both *A. thaliana* DCL1 and *D. melanogaster* Dicer-2 have a wide cleft within their helicase domains that can embrace double-stranded regions of primary pre-miRNA precursors (pri-miRNA) or pre-miRNA substrates. The helicase domains of insect Dicer-2 or plant DCL proteins can also bind long dsRNA substrates, like viral dsRNAs [28, 30]. Indeed, this feature of the helicase domains is characteristic of Dicers which are involved in protection against RNA viruses. These Dicers also display ATP-dependent translocation activity [2, 26, 27]. Conversely, the hDicer helicase domain has a much narrower cleft that cannot accommodate dsRNAs [9]. Actually, deletion of the hDicer helicase domain increases the efficiency of pre-siRNA substrate processing [57]. Moreover, a truncated Dicer variant lacking the HEL2i subdomain, a variant found in some mammalian cell lines, protects tissue stem cells from RNA viruses by dicing viral dsRNA [58]. It is believed that the helicase domains of vertebrate Dicers have lost their ability to recognize viral dsRNAs because vertebrates developed more complex immune systems [59].

In RNA binding assays using ssRNA substrates we noticed a HEL-concentration-dependent loss of the main substrate form (**Fig. 4a, Fig. 7a**, Supplementary **Fig. S10**). The most plausible reason for the observed phenomenon is the rearrangement of the structure of RNAs by the hDicer helicase, as demonstrated by comparative PAGE under native and denaturing conditions (**Fig. 7b**, Supplementary **Fig. S9**) and CD spectroscopy (**Fig. 8**) studies. Our observations are consistent with structural studies carried out for hDicer [11] and *Mus musculus* Dicer [60] in complex with pre-miRNA. Taking into consideration these data, we can assume that upon binding of pre-miRNA, Dicer, through its helicase domain, induces structural changes within the apical loop of pre-miRNA. These structural rearrangements may result in a better fit of the substrate into the catalytic site and, consequently, more precise substrate cleavage. Indeed, in cells producing a Dicer variant lacking the helicase domain, a non-homogeneous miRNA pool was observed [60]. Structural studies of the mouse pre-miRNA•Dicer complex also revealed a two-step mechanism of pre-miRNA cleavage by Dicer, as follows: (i) Dicer locked in the closed state recognizes the pre-miRNA and forms the pre-cleavage state and (ii) Dicer switches into an open state that allows loading of pre-miRNA into the catalytic site of Dicer [60]. The Dicer helicase domain plays a unique structural role in this process, i.e., it locks Dicer in the closed state, which facilitates pre-miRNA selection. Transition to the cleavage-competent open state is stimulated by the Dicer-binding partner TRBP [60]. These mechanistic changes linked to miRNA biogenesis are also very likely for hDicer [11]. Moreover, structural data collected for *D. melanogaster* Dicer-2 [56] and *A. thaliana* DCL1 [55] indicate that the proper orientation of the helicase domain is crucial for the specific substrate recognition and binding, as well as for further processing. We may further speculate that the helicase domain orchestrates substrate binding that is not followed by hDicer-mediated cleavage; indeed the passive binding of cellular RNAs has already been reported [35].

As mentioned above, due to steric constraints, the RIG-I subdomains of vertebrate Dicers are presumably not capable of recognizing viral dsRNAs to activate antiviral immunity [29]. This contrasts with the RIG-I subdomains that are present in insect [28, 61] and plant [62, 63] Dicer proteins. Nevertheless, recent literature indicates the important role of the hDicer helicase domain in triggering antiviral responses [34]. This previous study showed that during viral infection, the hDicer helicase binds to proteins that are involved in the antiviral response, such as DHX9, ADAR-1, and PKR kinase [34]. The results of the current study will similarly facilitate a better understanding of the role of hDicer in cellular processes extending beyond small RNA biogenesis pathways.

## Supplementary Information

The online version contains supplementary material.

## Statements and Declarations

### Author Contributions

.C., A.S., K.S., K.W. and A.U. planned and performed experiments: K.C. produced the HEL and hDicer_K70A protein preparations, conducted ATPase and binding assays, and BLI experiments; A.S. produced the hDicer_ΔHEL protein preparation, conducted the RNase assays; K.S. conducted CD and SAXS measurements; K.W. produced WT hDicer protein preparation; A.U. conducted BLI measurements; all authors analyzed the data and interpreted the results; A.K.K. conceived the studies, coordinated the research, supervised and provided advice; K.C. and A.S. wrote the draft of the manuscript; A.K.K. revised and edited the manuscript and was responsible for its final form. All authors have read and agreed to the published version of the manuscript.

### Data Availability

Additional data and datasets related to this paper are included as electronic supplementary material and are available from the corresponding author, A.K-K., upon reasonable request.

### Ethics approval and consent to participate

Not applicable.

### Consent for publication

Not applicable.

## Acknowledgments

We would like to thank Prof. Bryan R. Cullen for generously providing Dicer-deficient (NoDice) 293T human cell lines, Marta Wojnicka for providing the construct encoding WT hDicer and Paulina Krzemińska for a technical support in protein preparation. We also would like to thank Life Science Editors for editing services. We acknowledge the use of the EMBL SAXS beamline P12 at the Petra III storage ring of DESY, Hamburg, Germany.

