## Supplementary file for "The human Dicer helicase domain mediates ATP hydrolysis and RNA rearrangement"

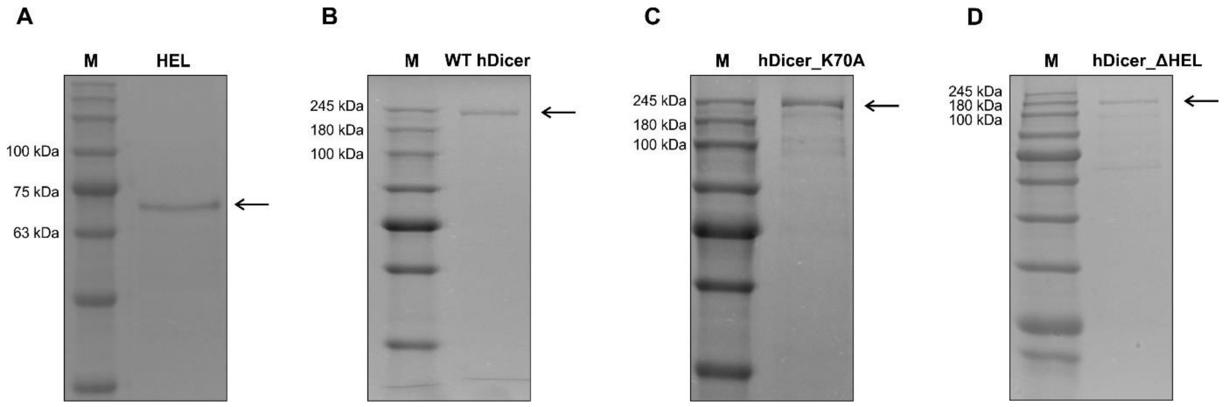


**Fig. S1** Protein preparations used in the study. SDS-PAGE gel showing: **a** the hDicer helicase domain preparation (HEL), 1 µg; **b** the wild-type hDicer protein preparation (WT hDicer), 0.5 µg; **c** the hDicer variant with a mutation in the Walker A motif (hDicer_K70A), 1 µg; and **d** the hDicer variant lacking the helicase domain (hDicer_ΔHEL), 0.5 µg. The respective proteins are indicated with arrows. HEL was produced in *Escherichia coli* as a fusion to the His6-tag. WT hDicer, hDicer_K70A and hDicer_ΔHEL were produced in HEK 293T NoDice cells [1] as fusions with the 3xFlag-tag. M indicates protein mass marker (*Perfect™ Tricolor Protein Ladder*, EURx).

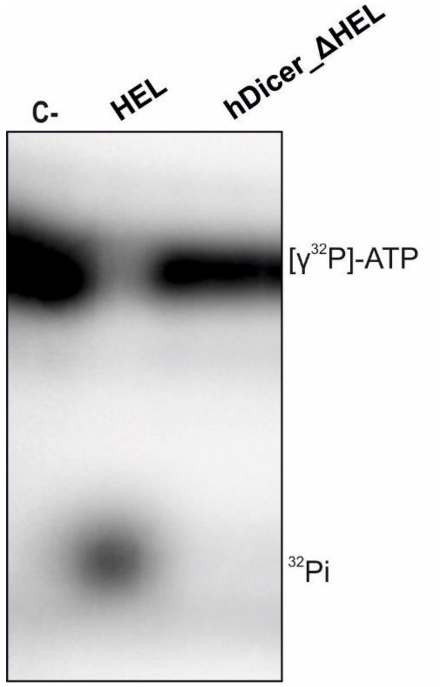


**Fig. S2** ATP hydrolysis assay with the hDicer helicase domain (HEL) and the hDicer variant lacking the helicase domain (hDicer_ΔHEL). The reactions involved: HEL (2 nM) or hDicer_ΔHEL (2 nM) and [$\gamma^{32}$P]-ATP (2 nM), and were incubated at 37 °C for 30 min. C- indicates a control sample with no protein and $^{32}$Pi indicates the product of [$\gamma^{32}$P]-ATP hydrolysis.

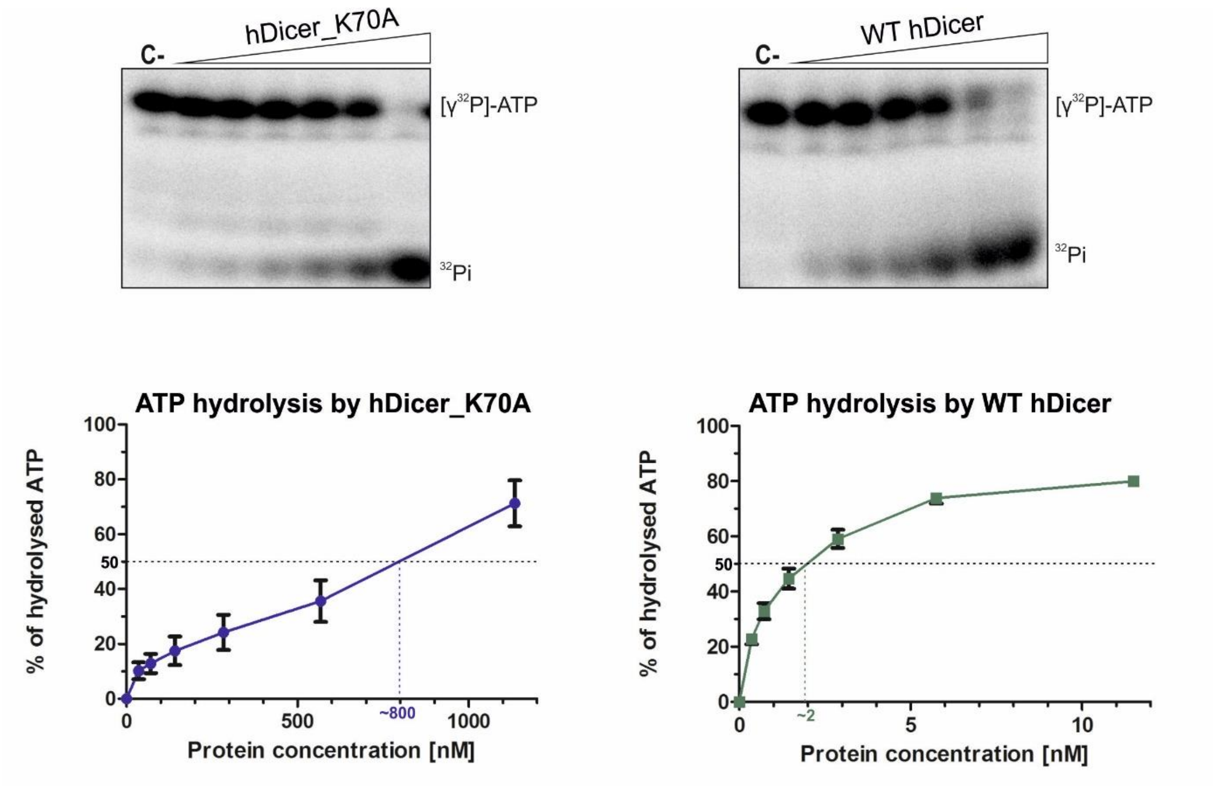


**Fig. S3** Protein concentration-dependent ATP hydrolysis of hDicer_K70A and WT hDicer. PAGE analysis of the reaction mixtures containing: $[\gamma^{32}P]$-ATP (2 nM) and hDicer_K70A (36, 72, 144, 288, 575, 1150 nM) or WT hDicer (0.36, 0.72, 1.44, 2.88, 5.75, 11.5 nM), reaction mixtures were incubated 30 min at 37 °C. C- indicates a control sample with no protein and $^{32}Pi$ indicates the product of $[\gamma^{32}P]$-ATP hydrolysis. Error bars represent standard deviations (SD) based on three separate experiments.

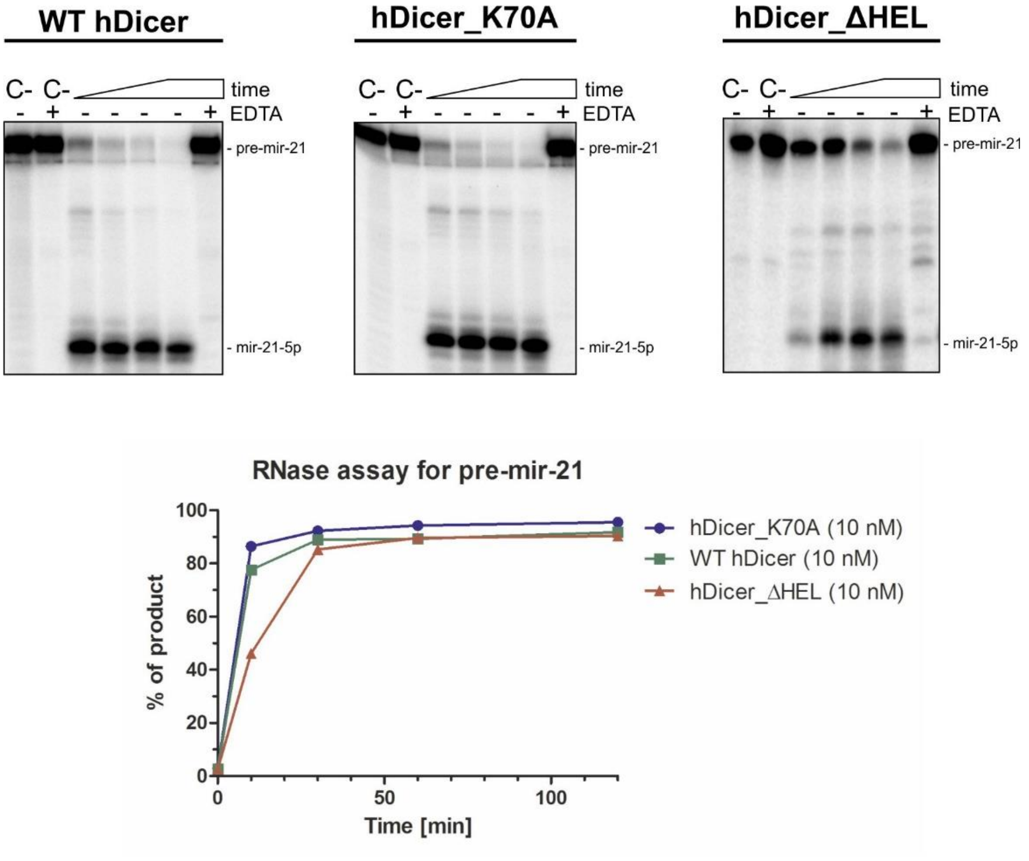


**Fig. S4** RNase activity assay involving WT hDicer, hDicer_K70A and hDicer_ΔHEL. PAGE analysis of the mixtures containing: pre-mir-21 (5 nM) and WT hDicer (10 nM) or hDicer_K70A (10 nM) or hDicer_ΔHEL (10 nM). Reaction mixtures were incubated at 37 °C for 10, 30, 60 and 120 min. C- indicates controls containing only the substrate in the reaction buffer. C+ indicates controls containing the substrate in the reaction buffer with the addition of the $Mg^{2+}$-chelating agent, EDTA (25 mM). +EDTA indicates samples where the reaction buffer was supplemented with 25 mM EDTA.

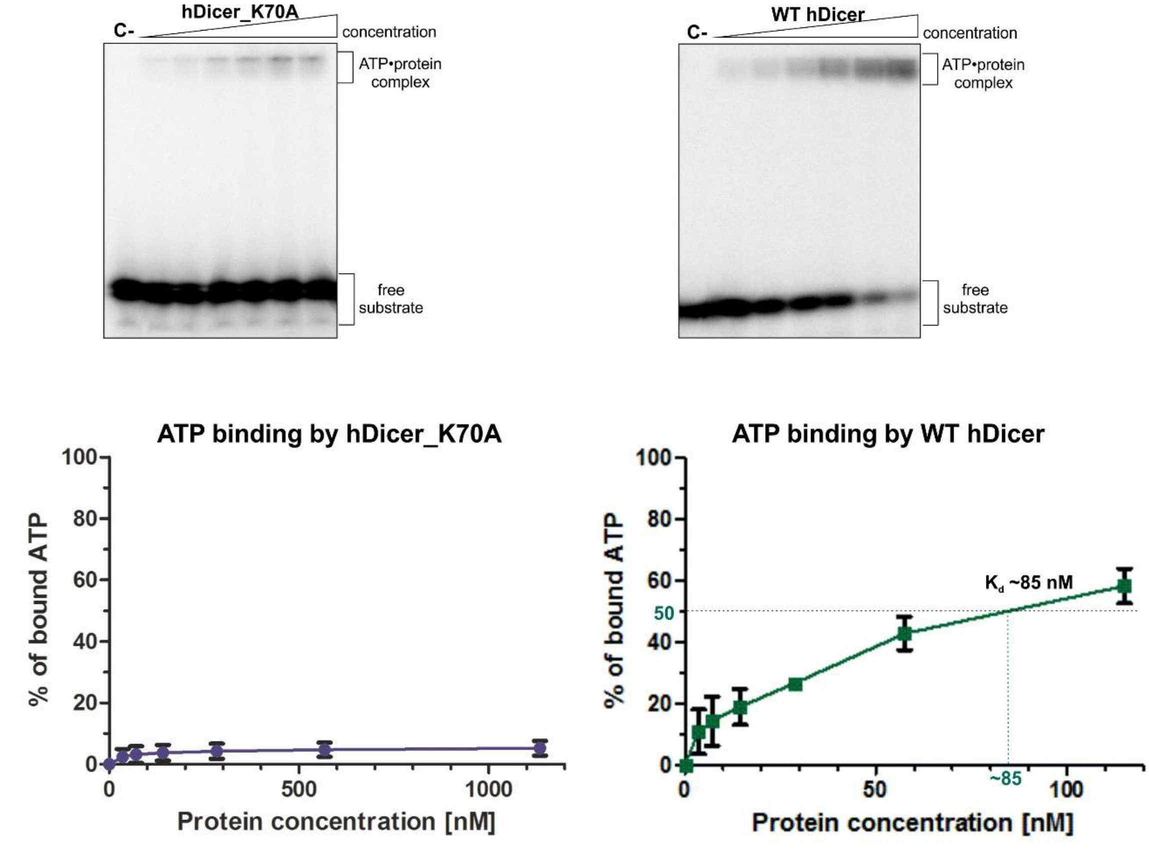


**Fig. S5** Protein concentration-dependent ATP binding of hDicer_K70A and WT hDicer. PAGE analysis of reaction mixtures containing: [$\gamma^{32}$P]-ATP (2 nM) and hDicer_K70A (36, 72, 144, 288, 575, 1150 nM) or WT hDicer (3.6, 7.2, 14.4, 28.8, 57.5, 115 nM), reaction mixtures were incubated 1 min at 4 °C. C- indicates a control sample with no protein. Error bars represent standard deviations (SD) based on three separate experiments.

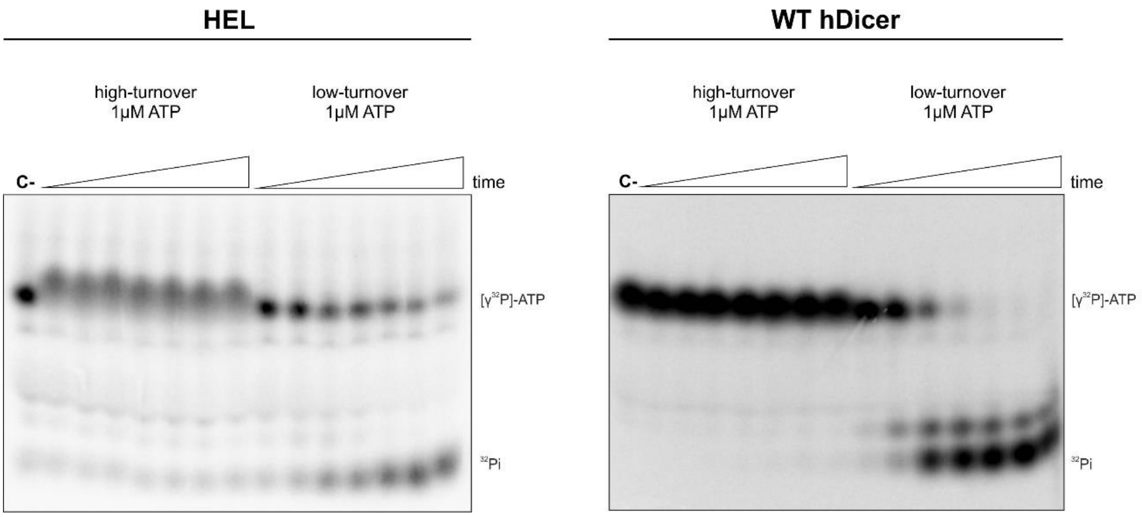


**Fig. S6** Time-course of ATP hydrolysis by HEL and WT hDicer under high-turnover and low-turnover conditions. PAGE analysis of reaction mixtures containing (high-turnover): 1 μM ATP with [$\gamma^{32}$P]-ATP (2 nM) spiked in to monitor hydrolysis and HEL (2 nM) or WT hDicer (2 nM), (low-turnover): [$\gamma^{32}$P]-ATP (2 nM) and HEL (2 nM) or WT hDicer (2 nM). Reaction mixtures were incubated at 37 °C for: 1, 5, 15, 30, 60, 90 and 120 min. C- indicates a control sample with no protein and $^{32}$Pi indicates the product of [$\gamma^{32}$P]-ATP hydrolysis.

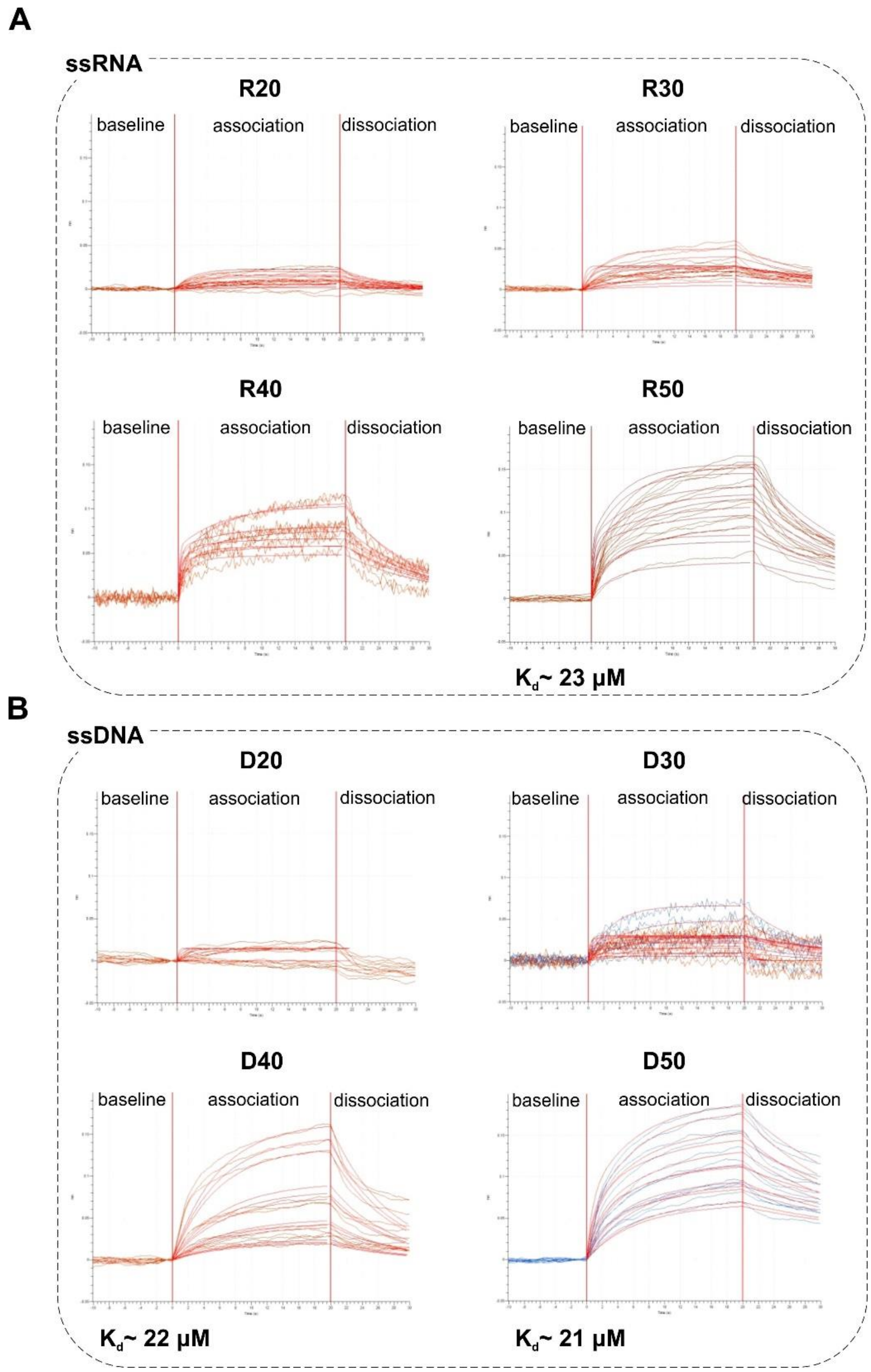


**Fig. S7** Nucleic acid binding activity of HEL measured using bio-layered interferometry (BLI). Binding curves obtained by BLI for **a** ssRNAs and **b** ssDNAs and HEL. In the experiment, HEL (1 μM) was incubated with increasing amounts of ssRNA (3.125, 6.25, 12.5, 25, 50 100 μM): R20, R30, R40, R50; or increasing amounts of ssDNA (3.125, 6.25, 12.5, 25, 50 100 μM): D20, D30, D40, D50. Measurements were carried out at 23 °C.

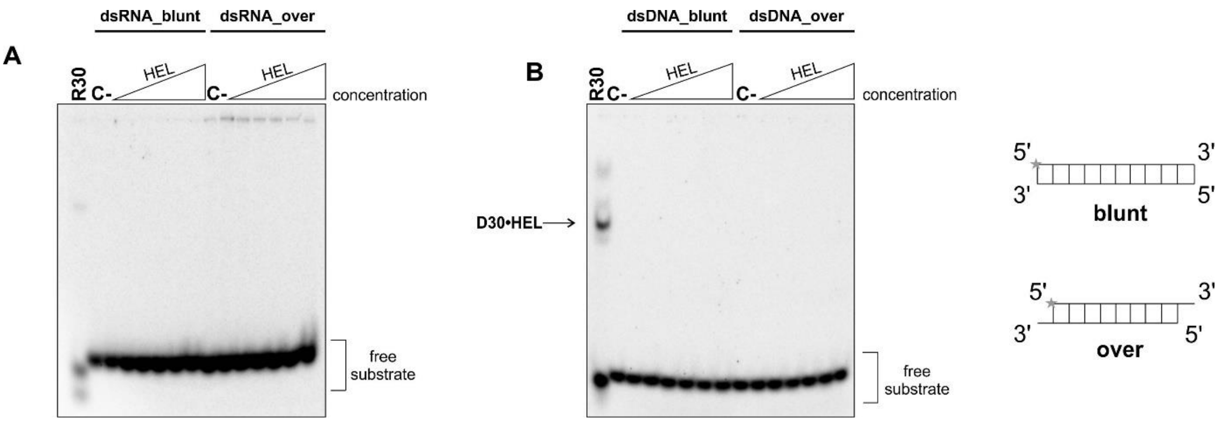


**Fig. S8** Double-stranded RNA (dsRNA) and double-stranded DNA (dsDNA) binding by HEL. **a** EMSA with HEL and $^{32}$P-labeled 32-bp RNA (dsRNA_blunt) (2.5 nM) and 30-bp RNA duplex with a 2-nt 3′ overhang on each end (dsRNA_over) (2.5 nM), R30 – a control sample: $^{32}$P-labeled R30 incubated with 95 µM HEL. **b** EMSA with HEL and $^{32}$P-labeled 32-bp DNA (dsDNA_blunt) (2.5 nM) and 30-bp DNA duplex with a 2-nt 3′ overhang on each end (dsRNA_over) (2.5 nM), D30 – a control sample: $^{32}$P-labeled D30 incubated with 95 µM HEL. Increasing amounts of HEL (2.97, 5.94, 11.86, 23.75, 47.5, 95 µM) are represented by a triangle. Reaction mixtures were incubated at room temperature for 15 min. C- indicates a control sample with no protein.

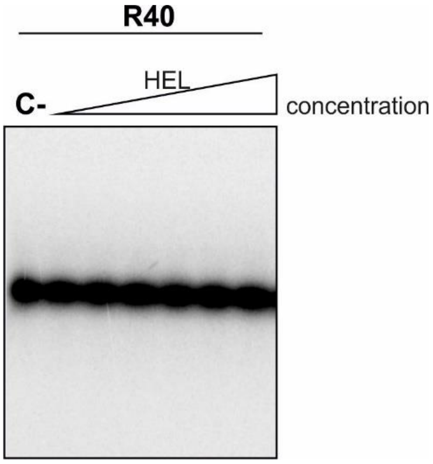


**Fig. S9** Denaturing PAGE analysis of 5′-$^{32}$P-labeled R40 (2.5 nM) incubated with increasing amounts of HEL (2.97, 5.94, 11.86, 23.75, 47.5, 95 µM). Reaction mixtures were incubated for 15 min at room temperature, then they were denatured at 95 °C in a loading buffer containing 10 M urea, and separated in an 8% denaturing (7 M urea) PAA gel. C- indicates a control sample with no protein.

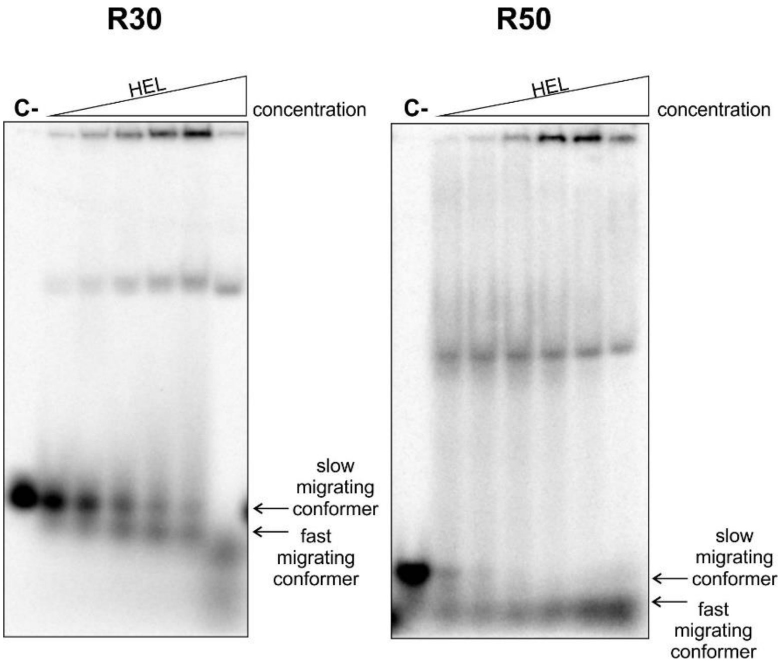


**Fig. S10** EMSA analysis of interactions between ssRNAs (R30, R50) and HEL. PAGE analysis of the mixtures containing: 5′-$^{32}$P-labeled R30 or R50 and increasing amounts of HEL (2.97, 5.94, 11.86, 23.75, 47.5, 95 µM). C- indicates a control sample with no protein.

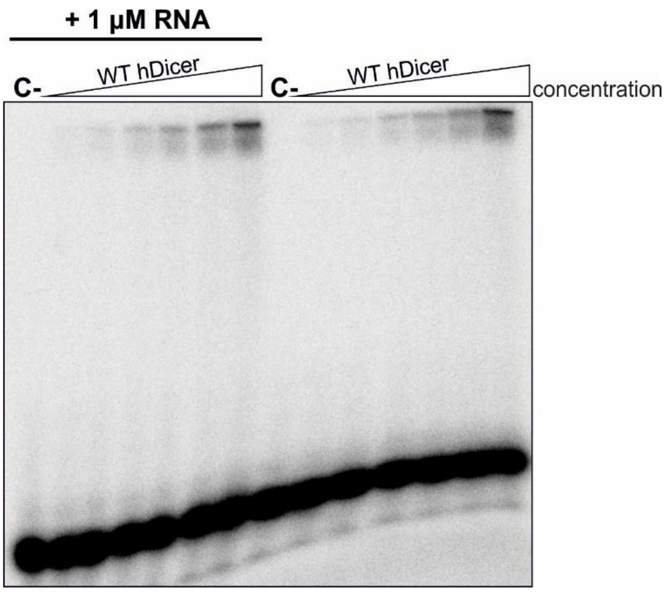


**Fig. S11** ATP binding by the wild-type hDicer (WT hDicer) with or without 1 µM non-labeled ssRNA. EMSA with [$\gamma^{32}$P]-ATP (2 nM) and increasing amounts of WT hDicer (0.45, 0.9, 1.8, 3.6, 7.2, 14.4 nM). Reaction mixtures were incubated at 4 °C for 1 min. C- indicates a control sample with no protein.

**Datasets related to the SAXS studies**

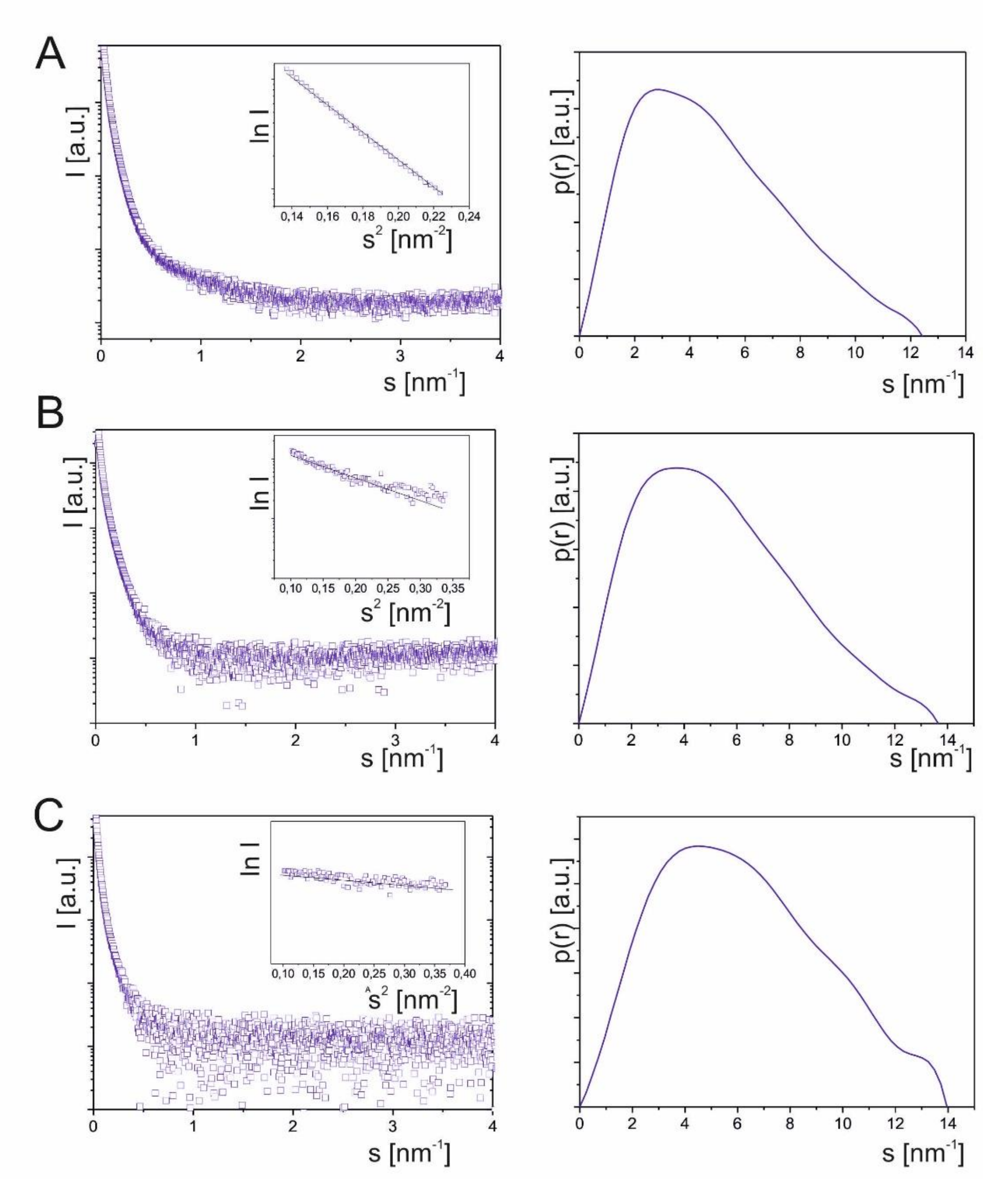


Experimental SAXS data. Plots present data for: **a** HEL, **b** HEL•pre-mir-21 complex, and **c** HEL•pre-mir-16-1 complex. Experimental SAXS curves (left) and Guinier plots (insert). A straight line in a Guinier region suggests that the system is essentially monodisperse. Pair distance distribution function, the *p(r)* function (right). The *p(r)* function is a histogram of all pairwise distances *r* between two scattering elements in the macromolecules weighted by their electron density contrast. The *p(r)* function is a representation of the shape of the molecule (or the complex of molecules) in real space. In the case of HEL and HEL complexes with pre-miRNAs, the *p(r)* function shows elongated shape of the analyzed molecules.

**Table S1** SAXS data collection and scattering-derived parameters.

| | HEL | pre-mir-21•HEL | pre-mir-16-1•HEL | pre-mir-21 | pre-mir-16-1 |
|---|---|---|---|---|---|
| Data collection | | | | | |
| Instrument | P12 beamline PETRA III storage ring | | | | |
| Wavelength (Å) | 1.24 | | | | |
| S range ($nm^{-1}$) | 0.0088-5 | | | | |
| Exposure time (s) | 1 | | | | |
| Temperature (K) | 293 | | | | |
| Structural parameters | | | | | |
| $I_0$ (arbitrary units ) [from P(r)] | 360.93 | 410.72 | 406.7 | - | - |
| $R_g$ (Å) [from p(r)] | 34.26 | 37.93 | 38.18 | - | - |
| $I_0$ (arbitrary units) (from Guinier region) | 360.93 | 410.75 | 405.6 | - | - |
| $R_g$ from crystal structure | 32.45 | - | - | 28.30 | 28.77 |
| Porod volume estimate ($Å^3$) | 84394 | 111284 | 115287 | - | - |
| Dry volume calculated from sequence ($Å^3$) | 92970 | 109840 | 111840 | 16870 | 18870 |
| Molecular-mass determination | | | | | |
| Contrast ($\Delta\rho x10^{10}$ $cm^{-2}$) | 3.047 | 3.047 | 3.047 | 3.047 | 3.047 |
| Molecular mass $M_w$ [from I(0)] (kDa) | 74689 | 94268 | 95605 | - | - |
| Molecular mass from sequence | 71462 | 89156 | 91202 | 17664 | 19740 |
| Software used | | | | | |
| Primary data reduction | PRIMUS | | | | |
| Data processing | PRIMUS | | | | |
| Quaternary structure modeling | SASREF | | | | |
| Computation of model intensities | CRYSOL | | | | |
| Three dimensional graphics representation | PyMOL | | | | |

**Table S2** Comparison of the theoretical and experimental thermodynamic parameters for R40 and the R40•HEL complex, calculated using Oligo Calc software (Biotools) [2] (theoretical) and circular dichroism (CD) spectroscopy (experimental).

| | Molecular mass of RNA [Da] | GC content [%] | $T_{mbasic}$ [°C] | $T_{msalt}$ [°C] | $T_{m\ exp.}$ [°C] | RlnK [cal/(°Kmol)] | ΔG [kcal/mol] | $\Delta G_{exp.}$ [kcal/mol] | ΔH [kcal/mol] | $\Delta H_{exp.}$ [kcal/mol] | ΔS [cal/(°Kmol)] | $\Delta S_{exp.}$ [cal/(°Kmol)] |
|---|---|---|---|---|---|---|---|---|---|---|---|---|
| R40 | 12741 | 50 | 69,4 | 79,9 | 35/75 | 33,404 | 58,6 | 56,7 | 347,5 | 336,03 | 915,5 | 885,28 |
| R40•HEL hDicer | 12741 | 50 | 69,4 | 79,9 | 60 | 33,404 | 58,6 | 50,2 | 347,5 | 297,81 | 915,5 | 784,58 |

$T_{mbasic}$ – basic melting temperature
$T_{msalt}$ – salt-adjusted melting temperature
$T_{m\ exp.}$ – experimental melting temperature
RlnK – ratio: R – gas constant (ok. 8,31 [$J*mol^{-1}*K^{-1}$]) and natural logarithm from K – equilibrium constant [mol/L]
ΔG – Gibbs energy
$\Delta G_{exp.}$ – experimental Gibbs energy
ΔH – theoretical enthalpy
$\Delta H_{exp.}$ – experimental enthalpy
ΔS – theoretical entropy
$\Delta S_{exp.}$ – experimental entropy
